# Autophagy inspects Rim4-mRNA interaction to safeguard programmed meiotic translation

**DOI:** 10.1101/2022.12.22.521672

**Authors:** Rudian Zhang, Wenzhi Feng, Suhong Qian, Shunjin Li, Fei Wang

## Abstract

Autophagy can degrade RNA, with an intricate preference for specific RNAs. However, the mechanism remains unclear. During yeast meiosis, autophagy is active, while a subset of transcripts needs to survive until their programmed translation during and at the end of meiotic divisions. Thus, the challenge here is how meiotic autophagy temporally spares specific mRNAs. Rim4, a meiosis-specific RNA binding protein (RBP), sequesters a specific set of mid-late meiotic transcripts during early meiosis to suppress premature translation. Recently, we reported that autophagy degrades Rim4, while the underlying mechanism and the fate of transcripts in complex with Rim4 remain uncharacterized. Here, we show that Rim4 utilizes a nuclear localization signal (NLS) to enter the nucleus to load its mRNA substrates before the nuclear export. Once in the cytoplasm, the active autophagy spares Rim4-mRNA. Using combined genetic, biochemical, and cell imaging approaches, we show that autophagy selectively degrades Rim4 during meiotic divisions in an Atg11-dependent manner upon Rim4-bound mRNAs released for translation; meanwhile, released mRNAs also become sensitive to autophagy. *In vitro*, purified Rim4 and its RRM-motif-containing variants, but not Rim4-mRNA complex, activate Atg1 kinase activity in meiotic cell lysates and in the immunoprecipitated Atg1 complex, suggesting that the conserved RRMs are involved in stimulating Atg1 and hence selective autophagy. These findings indicate that autophagy inspects Rim4-mRNA interaction to ensure meiotic stage-specific translation.

## Introduction

Selectivity is at the center of every cellular degradation pathway, including autophagy. This conserved eukaryotic pathway forms double-membraned vesicles (autophagosomes) and sends cytosolic contents to the lysosome (vacuole in yeast) for degradation. In *Saccharomyces cerevisiae* (budding yeast), autophagy is essential for meiosis and sporulation. Previous studies found that genetic deletion of essential autophagy-related genes (Atg) prevents meiotic DNA replication^1–5^. On the other hand, acute inhibition of autophagy at meiotic prophase I cause aberrant chromosome segregation and blocks meiosis exit^6,7^. Clearly, meiosis requires autophagy at multiple stages. Accordingly, a few autophagy substrates have been identified during meiosis, including the endoplasmic reticulum (ER), which is degraded by selective autophagy at the end of meiosis in an Atg41-dependent manner^8^; in addition, a negative meiosis regulator Ego4 is selectively degraded by autophagy at meiotic prophase I^1^. In line with these findings, we previously reported that autophagy degrades Rim4, a meiosis-specific RNA binding protein (RBP), during meiotic divisions^6^.

As the intermediate messenger, mRNA is typically needed for only a short period before its destruction. In cells, RBPs control the physiological and pathophysiological facets of the post-transcriptional processes of mRNAs, such as splicing, polyadenylation, and the editing, localization, stability, and translation^9–12^. Remarkably, Rim4 is essential for meiosis^13,14^; and, it is one of the most abundant meiotic RBPs that masterly controls mid-late meiotic translation^15–18^. Upon its expression, Rim4 sequesters mRNAs such as *CLB3* and *AMA1*, a B-type cyclin of Cdc28 (the yeast CDK1) and a meiosis-specific APC/C complex component, respectively, to prevent their premature translation^18^. At the end of meiosis I, the cellular level of Rim4 rapidly decreased due to proteasome-and autophagy-mediated degradation, allowing the translation of Rim4-sequestered mRNAs^6,16,19^. Therefore, through Rim4, autophagy is positioned to regulate the timing of meiotic translation.

Notably, the role of autophagy in Rim4 removal is irreplaceable; inhibition of autophagy during meiotic divisions prevents Rim4 clearance and hence *CLB3* translation^6^. While the cumulative phosphorylation of Rim4 by Ime2 stimulates proteasome-mediated Rim4 degradation^19^, it is unclear how autophagy degrades Rim4 in the context of meiotic translation. Unlike proteasomes, autophagy-mediated Rim4 degradation will potentially lead to the degradation of complex mRNAs. Intriguingly, it has been observed for a long time that autophagy can degrade RNAs across various organisms, including yeast. A recent study from Shiho Makino *et al*. shows that Rapamycin (TORC inhibitor)-stimulated autophagy prefers ribosome-associated mRNAs, indicating that autophagy can selectively degrades RNAs with an unknown mechanism^23^. Intriguingly, the downregulation of TORC activity is a feature of yeast meiosis, implying that meiotic mRNAs might be sensitive to selective autophagy. Therefore, the mechanism of autophagy-mediated Rim4 degradation centers on the fate of Rim4-sequestered mRNAs.

This study showed that the Rim4-mRNA complex is resistant to autophagy until the complex disassembly. Specifically, we identified an NLS that leads to Rim4 nuclear import. In the nucleus, Rim4 binds its mRNA substrates as a prerequisite for nuclear export. In the cytosol, the the Rim4-mRNA complex protects both Rim4 and mRNAs from autophagy. We show that upon mRNA released for translation during meiotic divisions, Rim4 is degraded by autophagy in an Atg11-dependent manner, involving Atg1 activation by RRM-containing Rim4. Meanwhile, mRNAs released from Rim4 are actively translated, while being sensitive to autophagy. Thus, we conclude that autophagy inspects Rim4-mRNA interaction to safeguard programmed meiotic translation.

## RESULTS

### A nuclear localization signal (NLS) promotes Rim4 nuclear import

Rim4 sequesters a subset of meiotic transcripts to suppress premature translation until its rapid degradation during the meiotic divisions^18^. Whereas RNA synthesis occurs in the nucleus, Rim4 suppresses translation in the cytoplasm^16,24^. To sequester its targets promptly, we propose that Rim4 approaches nascent mRNAs in the nucleus along the meiosis and sporulation. In this study, we adapted a β-estradiol inducible *GAL-NdT80* system to synchronize the entry into meiotic divisions to investigate Rim4 at specific meiotic stages (Fig. S1A)^25^. Particularly, without β-estradiol, the *GAL-NdT80* cells will be arrested at prophase I, a stage when Rim4 accumulates and suppresses translation^6,16,18,24^. With this system, we detected Rim4 (pRIM4:GFP-RIM4) signal by fluorescence microscopy (FM) in the nucleus of prophase I cells, marked by Nup49-mScarlet (*pNUP49:NUP49-mScarlet*), a nuclear pore complex (NPC) protein (Fig. 1A; 1B), or by RedDot™1, a membrane-permeable nuclear dye (Fig. S1B). As a control, the nuclear signal of Pab1 (*pPAB1:mScarlet-PAB1*), a Poly(A) binding protein that shuttles between the cytosol and nucleus^26–28^, is even lower than Rim4 (Fig. 1A). Next, we isolated nuclei from the prophase I cells. Like Pab1 (Pab1-FLAG), Rim4 (Rim4-V5) is detected in the isolated nuclei, while Por1 (mitochondria marker) and Hxk1 (cytoplasmic maker) are excluded, determined by immunoblotting (IB) (Fig. 1C). Thus, we conclude that Rim4 enters the nucleus before meiotic divisions.

**Figure 1.**
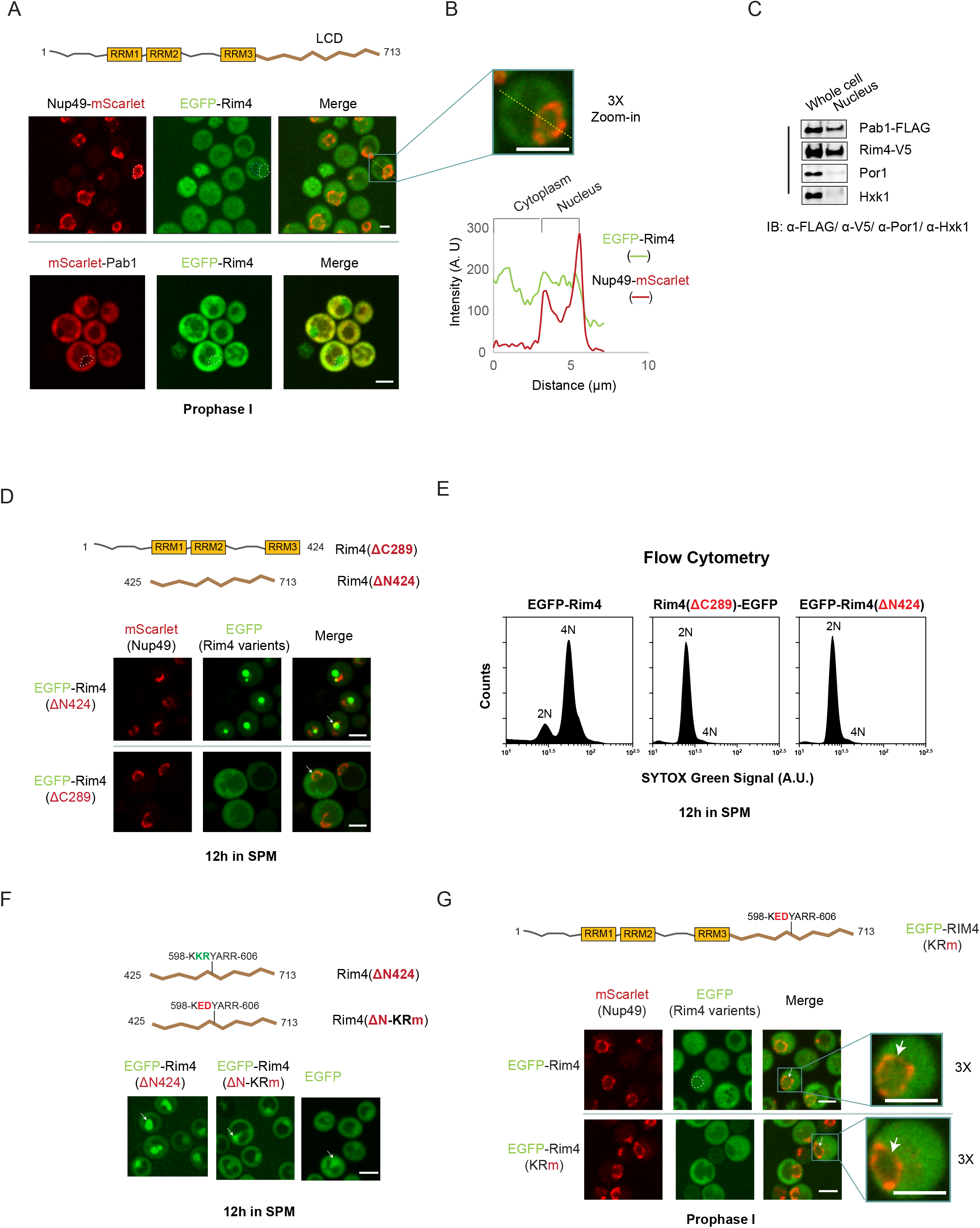
A nuclear localization signal (NLS) promotes Rim4 nuclear import. A. Top, the schematic of Rim4 protein, with three RNA recognition motifs (RRM) and a C-terminal low complexity domain (LCD). Middle and bottom, representative FM images of prophase I cells expressing mScarlet-Nup49/EGFP-Rim4 (middle) or mScarlet-Pab1/EGFP-Rim4 (bottom). The nuclear membrane is marked by mScarlet-Nup49 or by a dashed circle (white). A blue dashed circle marks the rough vacuole area. Details of the strains are in table 1 (strain list). The scale bar is 5 μm. B. Top, the 3x zoom-in of a cell from (A); Bottom, the intensity measurement of the EGFP (Rim4) and mScarlet (Nup49) fluorescent signals along indicated yellow dash line, as in the top panel. Rim4 resides in the nucleus and the cytoplasm. The peak of the mScarlet-Nup49 signal defines the nucleus-cytoplasm boundary. The scale bar is 5 μm. C. IB analysis of whole cell lysates and the isolated nucleus from prophase I cells expressing Pab1-Flag and Rim4-V5 under their endogenous promoter. Por1 is a mitochondrial outer membrane protein; Hxk1 is a cytoplasm marker; Pab1 shuttles between the nucleus and the cytoplasm. D. Top, the schematic of Rim4 variants. In Rim4(ΔN424), 424 residues are removed from the N-terminus; in Rim4(ΔC289), 289 residues are removed from the C-terminus. Middle and bottom, representative FM images of cells expressing mScarlet-Nup49 with EGFP-Rim4(ΔN424) (middle) or EGFP-Rim4(ΔC289) (bottom). The white arrow points to the nucleus. Time to collect cells for imaging: t = 12h in SPM. The scale bar is 5 μm. E. Graphs of DNA content quantification in meiotic cells expressing EGFP-Rim4, Rim4(ΔC289)-EGFP or EGFP-Rim4(ΔN424). After 12 hours in SPM, cells were fixed, stained with SYTOX Green, and analyzed by flow cytometry. F. Top, the schematic of the Rim4 variants. In Rim4(ΔN-KRm), K_600_R_601_ (green letter) of the Rim4(ΔN424) construct is replaced by E_600_D_601_; (red letter). Middle and bottom, representative FM images of cells expressing EGFP-Rim4(ΔN424), EGFP-Rim4(ΔN-KRm), or EGFP alone; all are driven by *RIM4* promoter. The white arrow points to the nucleus. The scale bar is 5 μm. G. Top, the schematic of Rim4 variants. Middle and bottom, representative FM images of prophase I cells expressing mScarlet-Nup49 with EGFP-Rim4 (middle) or EGFP-Rim4(KRm) (bottom). The white arrow points to the nucleus. Right, the 3x zoom-in view of selected cells. The Scale bar is 5 μm.

Rim4 is a ~80KD protein with a strong potential for self-assembly^16,17,19^; therefore, the nuclear import of Rim4 likely occurs via facilitated diffusion by binding to nuclear transport receptors (NTRs). Rim4 contains three RNA recognition motifs (RRMs) to its N-terminus (AA 1-424) and a C-terminal intrinsically disordered low complexity domain (LCD) (AA 425-713) (Fig 1D, top). By FM, we found that Rim4(ΔN424) (pRIM4: *GFP-Rim4[ΔN424]*), but not Rim4(ΔC289) (pRIM4: *GFP-Rim4[ΔC289]*), strongly prefers nuclear localization (Fig. 1D), suggesting that Rim4 carries a nuclear localization signal (NLS) in its C-terminal LCD region. This finding is consistent with the predicted nuclear localization of Rim4(ΔN424) by LocTree3 (Fig. S1C)^29^. Not surprisingly, both mutants failed to support meiotic DNA replication (Fig. 1E) and sporulation (Fig. S1D), indicating that functional Rim4 requires RRMs and the LCD. Next, we identified in Rim4’s LCD a seven-amino-acids sequence, 598-KKRYARR-606, similar to an NLS consensus sequence K-(K/R)-X-(K/R)^30,31^. Moreover, computational analysis using Alpha-fold predicts that this sequence and its flanking region are unstructured and thereby accessible to NTRs (Fig. S1E). Remarkably, mutating two residues at this site, i.e., K_600_E/R_601_D, sufficiently reduces the nuclear signal of Rim4(ΔN424) (Fig 1F) and Rim4 (Fig 1G). Thus, an NLS that contains K_600_R_601_ facilitates the nuclear import of Rim4.

### mRNA binding is a prerequisite for efficient Rim4 nuclear export

The nuclear retention of Rim4(ΔN424) (Fig. 1D), which lacks RNA-binding RRMs, implies that mRNA binding might be a prerequisite for Rim4 nuclear export; if so, it will be reminiscent of Pab1^26,27^. To directly examine this hypothesis, we constructed a Rim4 mutant, i.e., F139L/F349L (Rim4[FLm]), to destroy the conserved RNA binding sites in RRM-1 and RRM-3 motifs (Fig. 2A; S2A)^32^. As determined by electrophoretic mobility shift assay (EMSA), purified recombinant Rim4 (Rim4-6His), but not Rim4(FLm) (RIM4[FLm]-6His), robustly binds the *in vitro* synthesized 5’ UTR of *CLB3* mRNA (Fig. 2A), a model substrate of Rim4^18^. Like many RRM-containing RBPs, we found that Rim4 binds single-stranded DNAs (ssDNA) that match 120bps of the 5’ UTR of *CLB3* mRNA. The Rim4-ssDNA interaction was also reduced by the F139L/F349L mutation (Fig. 2A). Furthermore, the DNA replication and sporulation were abolished in Rim4(FLm) cells (Fig. 2B; 2C), confirming that Rim4(FLm) is defective in RNA binding, thereby dysfunctional.

**Figure 2.**
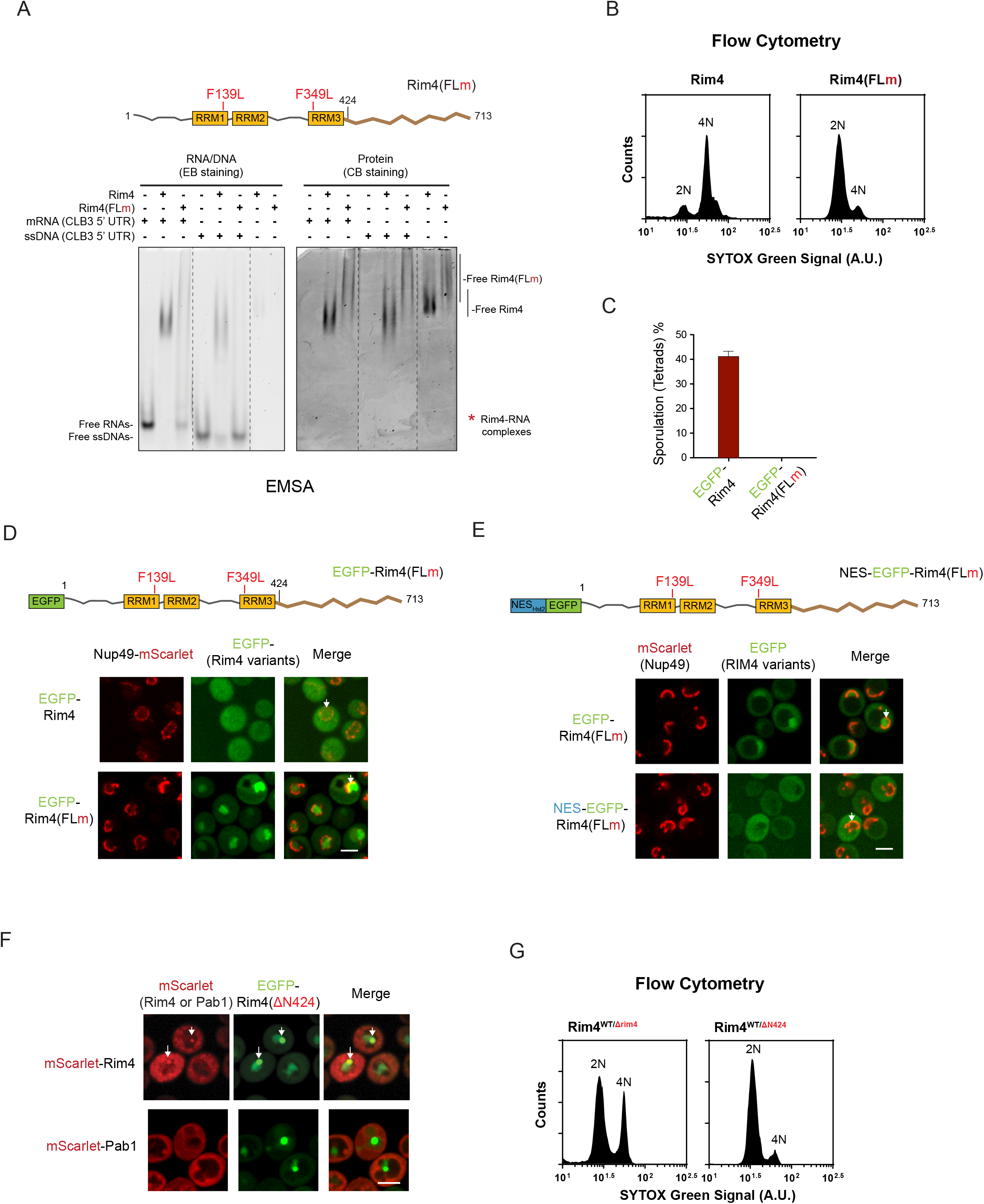
mRNA binding is a prerequisite for efficient Rim4 nuclear export. A) Top, the schematic of Rim4(FLm) with mutation site labeled. Bottom, Electrophoretic mobility shift analysis (EMSA) of *in vitro* synthesized mRNAs, or the ssDNA derived from the 5’ UTR of *CLB3*, binding Rim4 or Rim4(FLm). Left, Ethidium Bromide (EB) staining showing mobility shift of RNAs due to Rim4 or Rim4(FLm); right, Coomassie Brilliant blue (CB) staining showing mobility shift of indicated proteins due to mRNAs or ssDNAs. Red Asterisk marks the position of the co-migrated Rim4-mRNA or Rim4-ssDNA complex. B) Flow cytometry analysis of the DNA content (2N versus 4N) in the indicated Rim4 variants; cells were collected for measurement at 12 h in SPM before *NDT80* induction (*GAL-NDT80* system). C) The sporulation efficiency of indicated Rim4 variants, presented as the tetrads percentage; for each counting, n> 300 cells. D) FM analysis of cells (12h in SPM) shows nuclear retention of Rim4(FLm). Top, the schematic of Rim4(FLm); bottom, representative FM images of cells expressing Nup49-mScarlet with indicated EGFP-Rim4 variants. The white arrow points to the nucleus. The scale bar is 5 μm. E) Representative FM images of meiotic cells expressing Nup49-mScarlet and indicated GFP-Rim4 variants at prophase I. The white arrow points to the nucleus. The scale bar is 5 μm. F) FM analysis of prophase I cells show nuclear retention of Rim4(ΔN424) and its co-localization with Rim4, but not Pab1 in the nucleus. Shown are representative FM images of cells expressing EGFP-Rim4(ΔN424) with mScarlet-Rim4 (top) or mScarlet-Pab1 (bottom). The white arrow points to the nucleus. The scale bar is 5 μm. G) Flow cytometry analysis of indicated strains as described in (B).

Remarkably, when examined by FM, intracellular Rim4(FLm) (pRIM4:*EGFP-Rim4[FLm]*) mainly localizes in the nucleus, in sharp contrast to the wild type Rim4 (Fig. 2D), but reminiscent of Rim4(ΔN424) (Fig. 1D). This phenotype requires regular nuclear import of Rim4(FLm), because removal of LCD from Rim4(FLm) (Rim4[FLm-ΔC289]) effectively abolishes Rim4(FLm)’s nuclear retention (Fig. S2B). Moreover, a nuclear exit signal (NES) of Hst2^33^ fused to the N-terminus of protein rescued the nuclear retention phenotype in cells expressing Rim4(FLm) (NES-GFP-Rim4[FLm]) (Fig. 2E) or Rim4(ΔN424) (NES-GFP-Rim4[ΔN424]) (Fig. S2C), indicating that Rim4(FLm) and Rim4(ΔN424) are competent for nuclear export given an exit signal. We noticed that the nuclear retention of Rim4(FLm) is less penetrating than Rim4(ΔN424), possibly due to the residual RNA binding ability in Rim4(FLm) (Fig. 2A), because the RRM2 is still functional. Alternatively, Rim4 might carry a weak nuclear exit signal (NES) in its 1-424 region that enables slow mRNA-independent nuclear transport. Nonetheless, efficient Rim4 nuclear export relies on the Rim4-mRNA complex formation.

The coupling of Rim4-mRNA complex formation with Rim4’s nuclear export suggests that it is functionally relevant. Previous studies found that the intrinsically disordered LCD sequence mediates intracellular Rim4 self-assembly without interfering with Rim4-mRNA interaction^16^. Accordingly, we use Rim4(ΔN424) as a nuclear tether to disrupt regular Rim4 activity in the nucleus and the following Rim4 nuclear export. By FM, Rim4(ΔN424) specifically co-localizes with Rim4 in the nucleus but not Pab1, suggesting that Rim4(ΔN424) may interact explicitly with Rim4 in the nucleus (Fig. 2F). Strikingly, Rim4(ΔN424) exhibits a dominant negative effect on DNA replication (Fig. 2G) and sporulation (Fig. S2D), indicating that disrupted nuclear Rim4 activity impairs Rim4 function.

### The cytosolic Rim4-mRNA complex resists autophagy

Autophagy is active during yeast meiosis due to nutrient depletion, which is required for meiosis initiation and progression^1,6,8,34,35^. Thus, one immediate challenge a Rim4-mRNA complex faces in the cytosol is autophagy. Conceptually, Rim4-mRNA should survive autophagy to ensure the timely release and translation of Rim4-sequestered mRNAs during meiotic divisions^6,18^. As a probe, we developed a Nano-luciferase reporter fused with the 5’ and 3’ UTR of *CLB3*, a model substrate of Rim4 during meiotic divisions (Fig. S3A)^18^. We measured luciferase activity in meiotic cells over time and found that *CLB3 reporter* expression overlaps with cellular Rim4 removal, reminiscent of the Clb3 protein profile (Fig. 3A; S3B). In addition, inhibition of autophagy (Atg1-M102G[as] + 1NM-PP1 system) during meiotic divisions suppressed Clb3 translation, as previously reported^6^, and similarly reduced *CLB3* luciferase reporter activity (Fig. 3A; S3B). Thus, we hypothesize that autophagy degrades Rim4 after mRNAs are released, presumably to prevent Rim4 from binding the released mRNAs back. In other words, the Rim4-mRNA complex resists autophagy.

**Figure 3.**
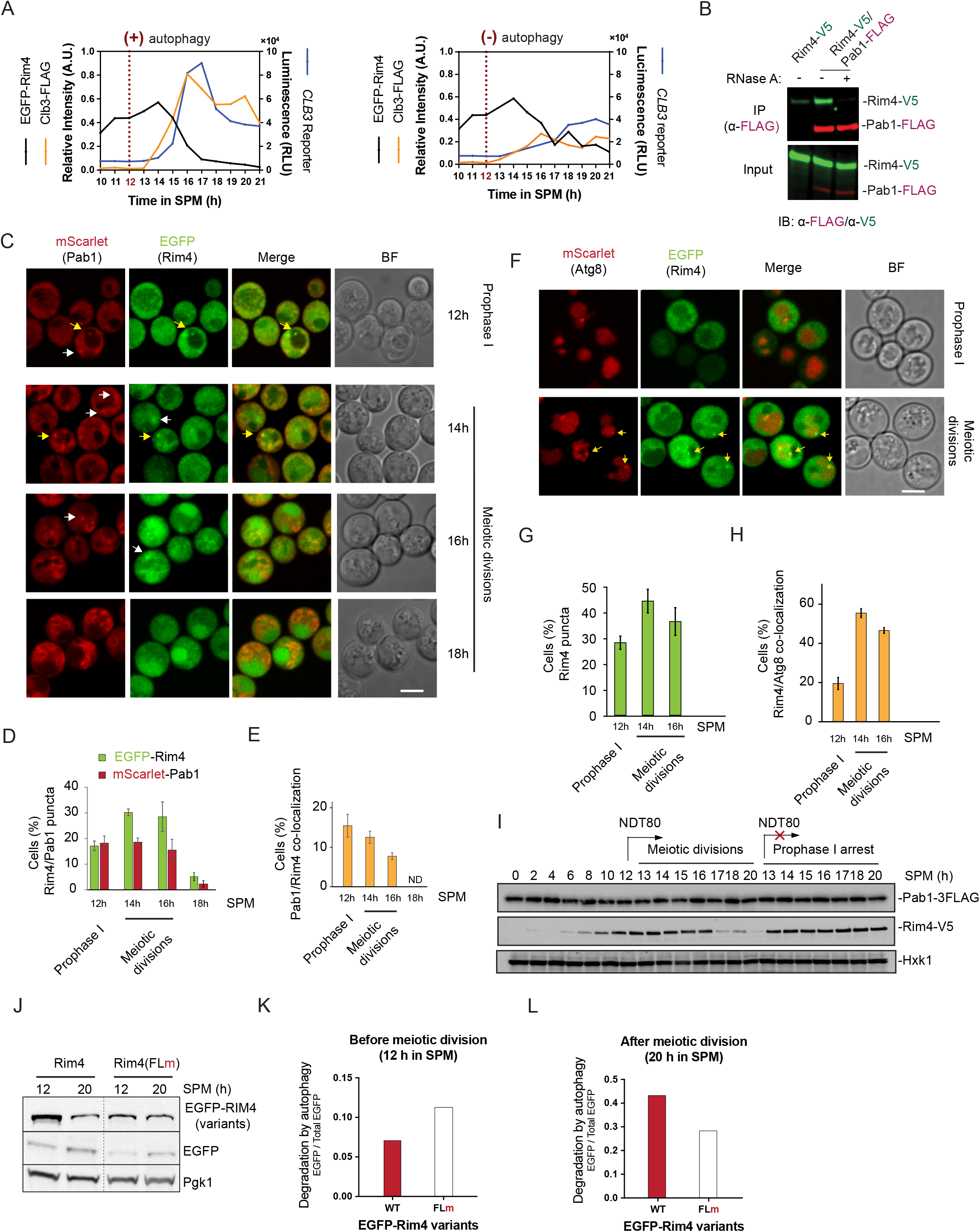
The cytosolic Rim4-mRNA complex resists autophagy. A) graph of cellular CLB3 reporter (nano-luciferase, described in Fig. S3A) luminescence signal (blue line), the EGFP-Rim4 (black line) and Clb3-FLAG (orange line) protein levels (determined by IB, Fig. S3B) during meiosis and sporulation. Left, autophagy is normal (- AI); right, autophagy is inhibited by 5 μM 1NM-PP1 (Atg1-as inhibitor) at 12h in SPM (+ AI). Clb3 protein synthesis correlates with Rim4 degradation. B) Cell extracts derived from meiotic prophase I cells that express Rim4-V5 and Pab1-FLAG were subjected to immunoprecipitation (IP) with magnetic beads-immobilized α-Flag antibody. The immunoprecipitated Pab1-3FLAG and its binding partners were eluted with 3xFlag peptide at 4 °C. Eluates (IP), and input samples were resolved by SDS-PAGE followed by IB with indicated antibodies. Applied RNase A was added to the cell lysate before IP. C-E) FM analysis of synchronized meiotic cells expressing EGFP-Rim4 and mScarlet-Pab1. Shown are (C) representative images; yellow arrow, co-localized mScarlet-Pab1 and EGFP-rim4 puncta; white arrow, mScarlet-Pab1 or EGFP-Rim4 puncta without co-localization; scale bars, 5μm; (D) percentage of cells with mScarlet-Pab1 or EGFP-Rim4 puncta; and (E) percentage of cells with Rim4/Pab1 co-localization. Only cells that carry EGFP-Rim4 puncta were analyzed here, ND, Non-detectable. (n≥300 cells, 3 replicate experiments, t-test). F-H) FM analysis of synchronized meiotic cells expressing EGFP-Rim4 and mScarlet-Atg8. Shown are (F) representative images; white arrow, co-localized mScarlet-Atg8 and EGFP-rim4 puncta; scale bars, 5μm; (G) percentage of cells with EGFP-Rim4 puncta; and (H) percentage of cells with Rim4/Atg8 co-localization. Only cells that carry EGFP-Rim4 puncta were analyzed here (n≥300 cells, 3 replicate experiments, t-test). I) IB of whole cell lysate with indicated antibodies showing Pab1-3Flag and Rim4-V5 protein level during meiosis and sporulation at indicated time points. Using an inducible GAL-NDT80 system, the protein level of Pab1-3Flag and Rim4-V5 were analyzed during normal meiotic divisions (+ β-estradiol at 12h in SPM), or during prophase I arrest (- β-estradiol). J-L) IB of cell lysates with indicated antibodies showing protein level of EGFP-Rim4 variants at prophase I (12h in SPM) and post-meiosis (20h in SPM). Pgk1, loading control. (K and L) The quantification of IB signals of (J) at indicated meiotic stages; free EGFP, and EGFP-Rim4 variants intensity was measured by BioRad Image Lab software, and the EGFP/Total EGFP signal ratio was calculated.

Rim4 binds mRNAs, which carry a poly(A) sequence that primarily interacts with Pab1. Remarkably, when we immunoprecipitated (IP) Pab1 (pPAB1: *PAB1-FLAG*) from synchronized meiotic cell lysates at prophase I, Rim4 was complexed with Pab1 in an RNA-dependent manner (Fig. 3B). Next, using Pab1 as an intracellular mRNA marker, we investigated whether autophagy degrades Rim4 in the absence of mRNAs by FM. Remarkably, upon entering meiotic divisions, the co-localization of Rim4 (EGFP-Rim4) puncta with Pab1 (mScarlet-Pab1) gradually disappeared over time (Fig. 3C; 3D; 3E), indicating that Rim4-sequestered mRNAs are released; while Pab1 still binds mRNAs to assist translation^28^. Concurrently, the Rim4 (EGFP-Rim4) puncta increased their co-localization with Atg8 (mScarlet-Atg8) (Fig. 3F; 3G; 3H), a marker of the forming double-membraned autophagosomes^36^, which send autophagic cargos to the vacuole (yeast lysosome) for degradation^37–40^. Notably, Pab1 does form cytosolic puncta but rarely colocalized with Atg8 (Fig. S3C; S3D). In addition, Pab1 remains stable when meiotic Rim4 degradation occurs, and the latter requires *NDT80*-induced meiotic divisions (Fig. 3I). Thus, Rim4 degradation by autophagy tightly correlates with the event of mRNA release during meiotic divisions.

Next, we examined the mRNA-binding defective Rim4 variants, GFP-Rim4[FLm] and GFP-Rim4[ΔN424]. Both mutants showed a weakened cytosolic protein signal than the wild type (Fig. 1D; 2D), supporting that mRNA-free cytosolic Rim4 is relatively unstable in the cytosol. Moreover, we found that EGFP-Rim4(FLm) are more sensitive to autophagy than EGFP-Rim4 before meiotic divisions (12h in SPM) when Rim4 was supposed to be stable, by IB analysis of the levels of the full-length protein and the vacuolar GFP procession resulted from autophagy-mediated degradation (Fig. 3J; 3K). The stability of Rim4 over Rim4(FLm) disappeared when Rim4 wild-type cells completed meiosis (20h in SPM) (Fig. 3J; 3L), in line with that Rim4 dissociates with mRNAs during meiotic divisions. Therefore, we conclude that the nuclear Rim4-mRNA assembly protects Rim4 from autophagy in the cytosol and presumably also protects Rim4-sequestered mRNAs, which we will examine later.

### Selective autophagy temporally degrades Rim4

Autophagy-mediated degradation could be non-selective (bulk) or selective. Since autophagy can discriminate Rim4 from the Rim4-mRNA complex, we examined whether Rim4 is degraded by selective autophagy. Earlier, we showed that Rim4 (EGFP-Rim4) was recruited to Atg8-positive (mScarlet-Atg8) autophagosomes; the percentage of Rim4 (GFP-Rim4) puncta colocalization with Atg8 increased from ~17% (prophase I) to ~56% (meiotic divisions) (Fig. 3H), indicating that autophagy mainly degrades Rim4 during meiotic divisions. Next, we constructed a strain expressing both EGFP-Rim4 and mCherry-GFP, each driven by *RIM4* promotor for equivalent expression (Fig. 4A), considering mCherry-GFP as a cargo of bulk autophagy. In this strain, based on the result of IB (α-EGFP), Rim4 (EGFP-Rim4) is more stable than mCherry (EGFP-mCherry) until *NDT80*-induced meiotic divisions when the relative stability of Rim4 to mCherry abruptly decreased (Fig. 4A; 4B). Moreover, the acute inhibition of autophagy (Atg1-as + 1NM-PP1 system)^6^ during meiotic divisions increased the relative GFP-Rim4 stability to mCherry-GFP (Fig. 4A; 4B). These results indicate that autophagy selects Rim4 over mCherry for degradation during the meiotic divisions. Next, we examined the relative stability of Rim4 over Ape1, an aminopeptidase deemed a selective substrate of the yeast Cytosol-to-Vacuole Targeting pathway (CVT), which is one type of selective autophagy. Similarly, we found that autophagy prefers degrading Rim4 over Ape1 during, but not before, meiotic divisions (Fig. S4A, S4B), suggesting that Rim4 is strongly preferred by selective autophagy during meiotic divisions.

**Figure 4.**
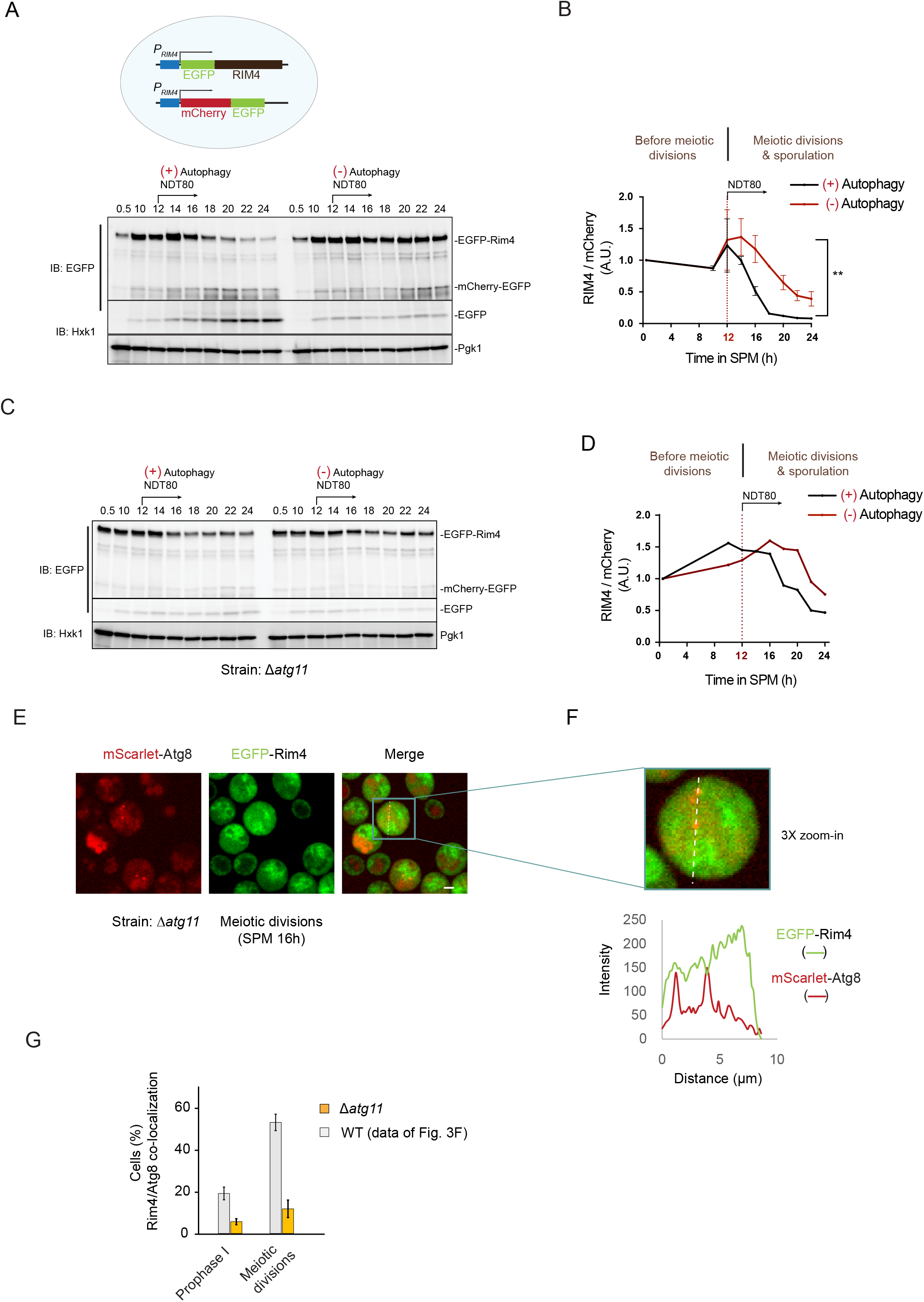
Selective autophagy temporally degrades Rim4. A and B) IB analysis with indicated antibodies (A) of cell lysate and quantification (B) of signal intensity showing EGFP-Rim4, EGFP-mCherry, and free GFP protein level during sporulation at indicated time points in SPM. (A) Top, the schematic of a cell harboring one allele of EGFP-Rim4 and one allele of EGFP-mCherry, under *the RIM4* promoter. Bottom, representative IB images. Pgk1, loading control. NDT80 expression was induced at 12h in SPM. (+) Autophagy, mock treatment; (-) autophagy, 5μM 1NM-PP1 treatment at 12h in SPM. (B) Graph showing ratio of EGFP-Rim4/EGFP-mCherry IB signal (2 replicate experiments, paired t-test). AU, arbitrary units. C and D) IB analysis as described in (A) and (B), except that *ATG11* was deleted from the genome (*Δatg11). ATG11* deletion increases relative Rim4 (Rim4/mCherry) stability, especially during meiotic divisions. E and F) Representative FM images of synchronized meiotic *Δatg11* cells (t = 16 h in SPM) that express mScarlet-Atg8 and EGFP-Rim4 (E), and intracellular fluorescent signal intensity analysis of mScarlet-Atg8/GFP-Rim4 (F). Scale bars, 5μm. (F), Fluorescence intensities (bottom) along the yellow dash line in a cell (top). G) Graph of mScarlet-Atg8/EGFP-Rim4 puncta co-localization analysis during prophase I and the meiotic divisions in *Δatg11* cells. *ATG11* wild-type data is from Fig. 3C as a comparison.

The above observations led us to examine the stability of meiotic Rim4 over time in *Δatg11* cells. In yeast, selective autophagy generally requires Atg11, a scaffolding protein that functions at phagophore assembly sites (PAS), where the autophagy machinery assembles upon autophagy induction. Remarkably, Rim4 becomes more stable in the absence of Atg11, especially during meiotic divisions (Fig. 4C; 4D). Moreover, during meiotic divisions, only ~12% of Rim4 puncta co-localize with Atg8 in *Δatg11* cells (Fig. 4E; 4F; 4G), decreased from ~56% as in WT cells (Fig. 3H). Notably, Atg8/Rim4 co-localization was also reduced to ~6% in *Δatg11* cells before meiotic divisions (prophase I) (Fig. 4G), as compared to ~17% in WT cells (Fig. 3H). Collectively, our data demonstrate that Atg11-dependent selective autophagy mediates Rim4 degradation. Importantly, inefficient Rim4 degradation in *Δatg11* cells correlates with reduced sporulation efficiency (Fig. S4C), suggesting that the mechanism that ensures rapid autophagic degradation of selected substrates, e.g., Rim4, is important for meiosis and sporulation.

### The Rim4, but not Rim4-mRNA, stimulates Ata1 activity in vitro

Next, we asked how autophagy discriminates Rim4 from the Rim4-mRNA complex. The ULK1(Atg1) is the master protein kinase of autophagy machinery required for autophagy initiation. The interaction of substrates with ULK1 (Atg1) kinase complex through a receptor has recently been reported by several groups and is an emerging concept in selective autophagy^41–44^. In addition, a substrate of selective autophagy can activate Atg1^41^. Therefore, we examined whether Rim4 activates Atg1 (Atg1-as) *in vitro* (Fig. 5A). Specifically, we prepared cell lysates from *atg1-as* cells arrested at meiotic prophase I due to lack of *NDT80* expression, a stage before Rim4 removal, and measured Atg1-as kinase activity corresponding to the supplementation of purified recombinant Rim4 protein (Rim4-6His) (Fig. S5A). Remarkably, recombinant Rim4 stimulates Atg1-as kinase activity, best when [Rim4] is ~0.3 μM (Fig. 5B), within the range of endogenous Rim4 cellular level during meiosis (0.2 to 1.2 μM, unpublished). Strikingly, the yeast total RNAs at the concentration of ~42 ng/μL abolished Rim4-stimulated Atg1-as kinase activity *in vitro* (Fig. 5B). Therefore, mRNA negatively affects Atg1 activation by Rim4.

**Figure 5.**
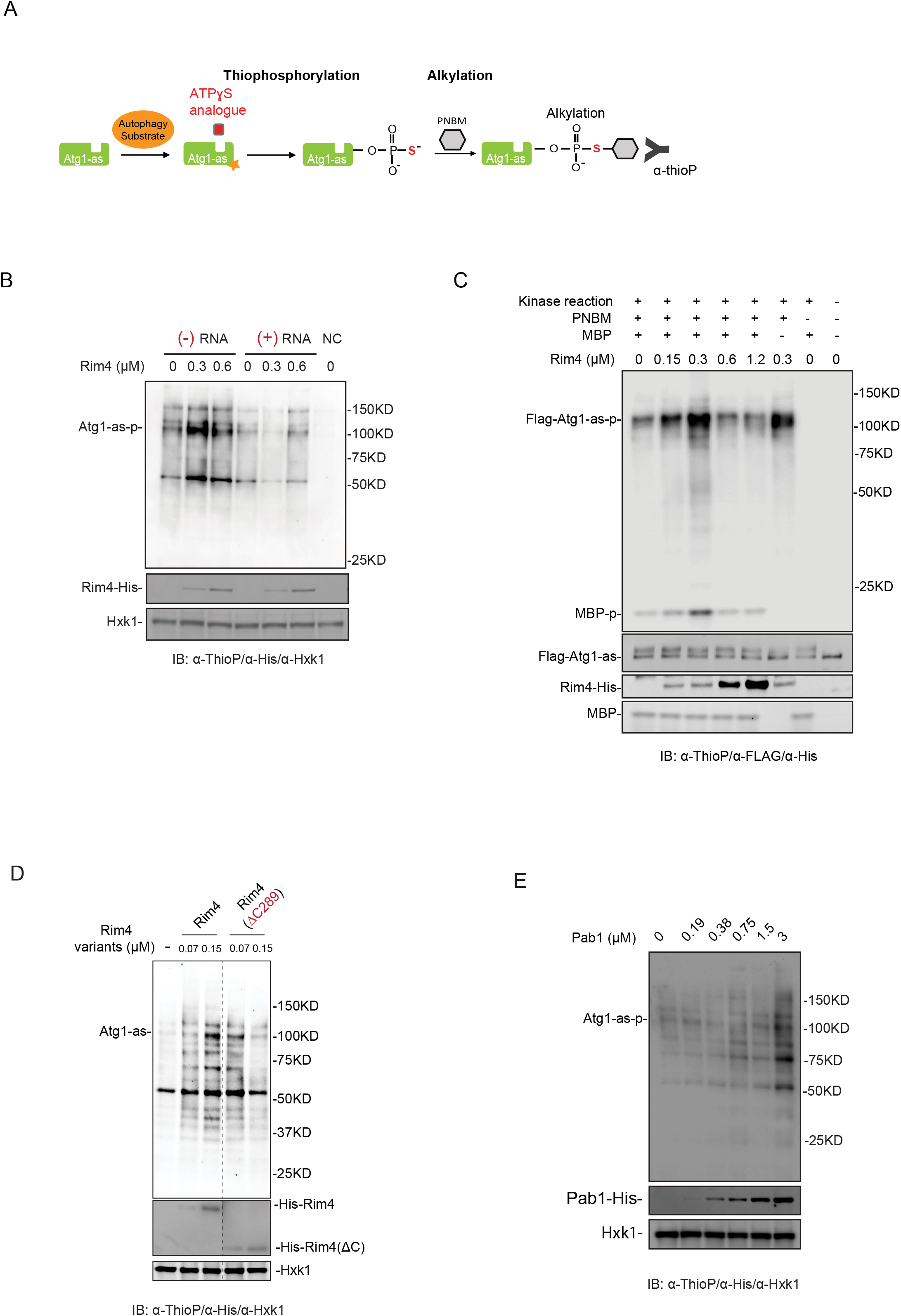
The Rim4, but not Rim4-mRNA, stimulates Atg1 activity in vitro. A) Schematic of the chemical-genetic strategy for monitoring Atg1-as (analog-sensitive Atg1, Atg1-M102G) kinase activation by a selective autophagy *substrate in vitro*. Atg1-as thiophosphorylates its substrates, including itself with a bulky ATPγS analog (N6-PhEt-ATP-γ-S). Thiophosphorylated substrates of Atg1-as, including Atg1-as itself (Atg1 auto-phosphorylation), can be alkylated with para-nitrobenzyl mesylate (PNBM) and then detected by IB using anti-thiophosphate ester (α-thioP) antibodies. B) Atg1-as cells under synchronized meiosis (Prophase I, SPM 12h) were collected; Atg1-as activity in prepared cell lysates was assayed as in (A). Before the Atg1-as kinase assay, cell lysates were supplemented with the indicated amount of recombinant Rim4-6xHis or premixed Rim4-6xHis/RNAs (final [RNA]=41.7 ng/μL). After the reaction, the whole-cell lysates were subjected to IB with indicated antibodies. Atg1-as auto-phosphorylation (Atg1-as-p) indicates Atg1-as kinase activity. Hxk1, loading control; NC, no supplemented ATP in cell lysate. C) FLAG-Atg1-as protein was affinity-purified (IP: α-FLAG) from extracts of meiotic cells arrested at prophase I, and then supplemented with indicated recombinant Rim4-6xHis ([Rim4] from 0.15 μM to 1.2 μM), followed by Atg1-as kinase assay as diagrammed in (A). Shown are IB of FLAG-Atg1-as complex after kinase assay with indicated antibodies. MBP, Myelin basic protein, is generic *in vitro* kinase substrate. The Atg1-as mediated thiophosphorylation of MBP and Atg1-as auto-phosphorylation agree on Rim4 stimulating Atg1 activity. D and E) In vitro Atg1-as assay as in (B), except that recombinant Rim4-6xHis and Rim4(ΔC289)-6xHis (D), or Pab1-6xHis (E), was supplemented to the Atg1-as cell lysates before conducting the kinase assay.

In line with Rim4 activates Atg1 to be degraded, inhibition of Atg1 kinase activity (Atg1-as and 1NM-PP1 system) during meiotic divisions results in reduced cytosolic co-localization of Rim4 puncta with the autophagosome marker Atg8 (Fig. S5B, S5C). Next, we tested the effect of Rim4 on the immunoprecipitated (IP) Atg1 (FLAG-Atg1-as) complex. Like that in the cell lysates, Rim4 stimulates affinity-purified Atg1 kinase activity in a dose-dependent manner (Fig. 5C, 0.3 μM), indicating that the intrinsic Rim4 property is sufficient to stimulate Atg1 in the Atg1 complex. This finding led us to examine the role of Rim4 RRM motifs in Atg1 activation. Accordingly, we purified recombinant Rim4(ΔC289) protein and Pab1, which contain RRM domains like Rim4. *In vitro*, both proteins activate Atg1-as in the cell lysates, albeit with different optimal concentrations (Fig. 5D, 5E). Thus, we speculate that autophagy recognizes Rim4 at least partially via RRMs, which can be masked by mRNA binding.

### Timina of Rim4 degradation is strictly programmed to support meiosis and sporulation

Previous studies have demonstrated that defective Rim4 removal during meiotic divisions causes defects in meiosis and sporulation^16,18^. However, the effect of premature Rim4 degradation on meiosis and sporulation was not studied, except that Rim4 is essential for meiotic DNA replication based on genetic studies^13,14^. Therefore, we engineered N-Degron tagged Rim4 (N-Deg-Rim4-V5) to allow β-estradiol-triggered Rim4 removal by proteasomes while monitoring meiotic DNA replication and sporulation (Fig. 6A). As a proof of concept, the N-deg-Rim4-v5 protein exhibited a WT-like meiotic profile (Fig. S6A) and sporulation (Fig. S6B), and its level diminished after ~2h of β-estradiol triggered degradation (Fig. S6A). Strikingly, premature Rim4 degradation, i.e., adding β-estradiol before meiosis-scheduled Rim4 clearance, leads to reduced sporulation efficiency (Fig. 6B); the earlier was N-deg-Rim4-v5 degraded, the more severer was the sporulation defect (Fig. 6B). This result indicates that premature degradation of Rim4, and hence the Rim4-mRNA complex, is harmful to sporulation.

**Figure 6.**
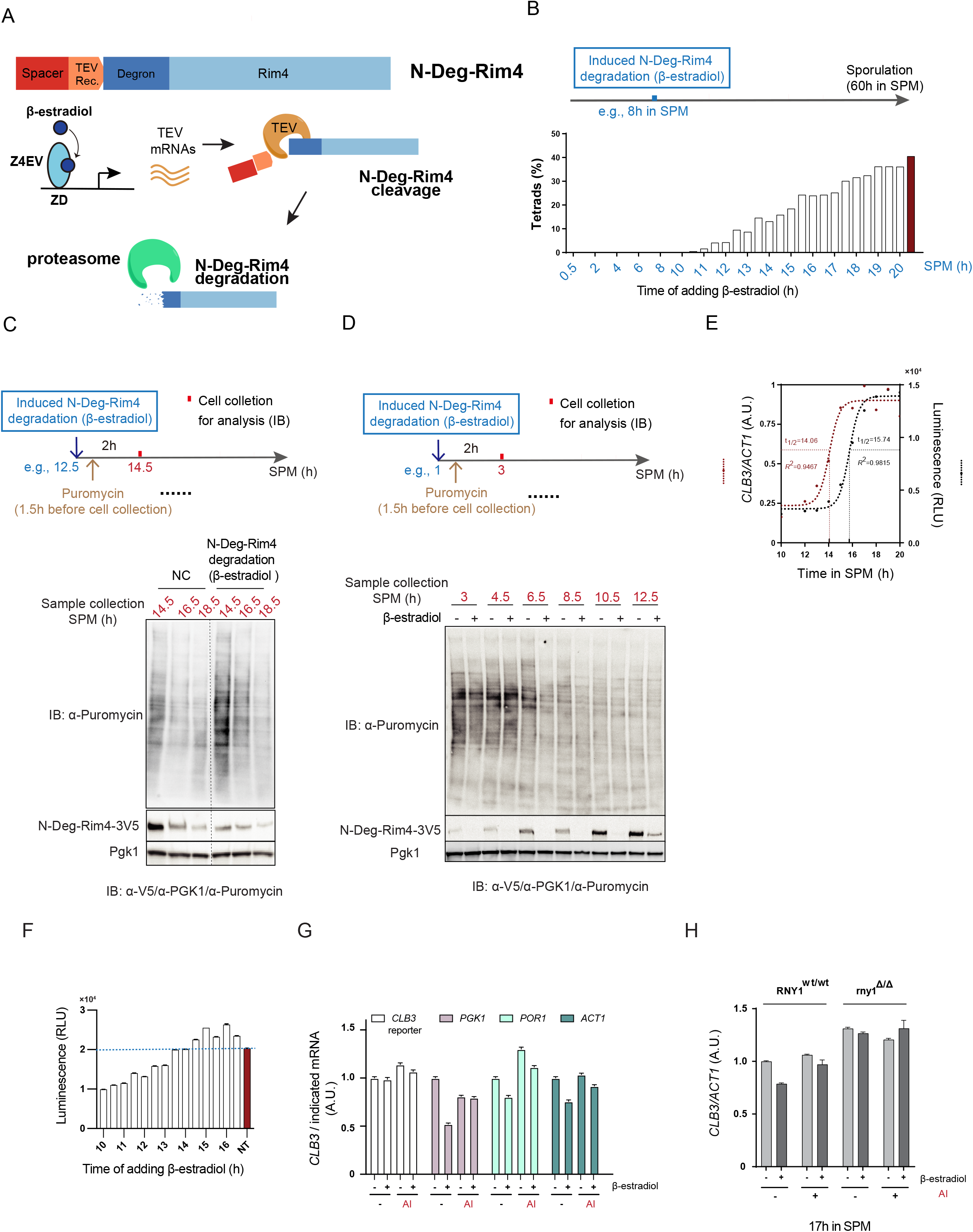
Timing of Rim4 degradation is strictly programmed to support meiosis and sporulation. A) Inducible Rim4 degradation by the proteasome. Top, a construct of N-Deg-Rim4 comprises a TEV recognition site to Rim4’s N-terminus that allows TEV mediated cleavage; bottom, β-estradiol-induced expression of TEV (*pZEV: TEV*), which cleaves N-Deg-Rim4, resulting in the exposure of an N-degron for stimulated proteasome-mediated Rim4 degradation. B) Sporulation (tetrad formation) efficiency plotted to the timing of β-estradiol-induced N-Deg-Rim4 degradation (n>300 cells for each time point). C and D) Active protein synthesis by ribosomes was measured after 2.5h of β-estradiol-induced N-Deg-Rim4 degradation. 1 μM β-estradiol was added to SPM before meiotic divisions (C) or during meiotic divisions (D) at the indicated time. To label protein synthesis, 10 μg/mL Puromycin was added to the culture 30 min after applying β-estradiol or mock treatment and incubated for two hours; then, cells were harvested for IB analysis using indicated antibodies. E) Intracellular *CLB3* mRNA level (RT-qPCR), and *CLB3* reporter translation (Luciferase assay), were plotted to the time of cells in SPM. Then, a sigmoidal regression was fitted to both set of data; and the t_1/2_ were calculated, showing a ~2h gap between *CLB3* transcription and translation. The CLB3 mRNA level at the peak value was normalized to 1. F) Graph of *CLB3* luciferase reporter activity in N-Deg-Rim4 cells at 17h in SPM. 1μM β-estradiol at indicated time points was applied to the SPM to trigger N-Deg-Rim4 degradation. NT, cells were not treated with β-estradiol. G) Graph of *CLB3* mRNA levels (ΔCt analysis) in cells, relative to indicated *mRNAs*. If applied, the β-estradiol (inducing N-Deg-Rim4 degradation) and/or autophagy inhibitor 1NM-PP1 (AI) were added at 15h in SPM. After 2h of treatment in SPM (12h in SPM), cells were collected for RNA extraction, cDNA reverse-transcription, and qPCR. The relative CLB3 mRNA levels without any treatment (β-estradiol -, AI -) are set as 1. H) Graph of *CLB3* mRNA levels (ΔCt analysis) in cells at 17h in SPM. Tested strains include *RNY1* WT (RNY1^WT/WT^) and *RNY1* deletion (rny1^Δ/Δ^). If needed, the β-estradiol and/or autophagy inhibitor 1NM-PP1 (AI) were applied at 15h in SPM. After 2h of treatment in SPM, cells were collected for RNA extraction, cDNA reverse-transcription, and qPCR. The CLB3 mRNA level from the RNY1WT strains without any treatment (β-estradiol -, AI -) was normalized to 1.

Next, we examined the effect of conditional removal of N-Deg-Rim4-v5 on translation at different meiotic stages, using a Puromycin labeling assay. Puromycin is a tyrosyl-tRNA mimic that blocks translation by labeling and releasing elongating polypeptide chains from translating ribosomes. Afterward, the Puromycin-labeled nascent peptides in cell lysates can be detected by IB (α-Puromycin). Using this assay, we found that β-estradiol triggered N-Deg-Rim4-v5 degradation stimulates translation during meiotic divisions, consistent with the role of Rim4 as a translation suppressor (Fig. 6C). Strikingly, stimulated translation by N-Deg-Rim4-V5 removal is restricted to ~12.5h to ~16.5h in SPM (Fig. 6C; 6D), specifically overlapping with the window of Rim4-suppressed *CLB3* translation (~12h to ~17h in SPM) (Fig. 6E). This result demonstrates that the Rim4 degradation to stimulate mid-late meiotic translation is strictly assigned to the meiotic divisions.

Notably, upon induction, proteasome removes N-Deg-Rim4-V5 regardless of its state of mRNA binding, while autophagy will spare the N-deg-Rim4-mRNA complex, as we demonstrated in this study. In line with that inducing N-Deg-Rim4-V5 removal during meiotic divisions substantially enhanced translation (Fig. 6C), increased level of CLB3 Luciferase reporter was observed at the end (17h in SPM) if N-Deg-Rim4 degradation was induced between 14.5h and 17.5h in SPM (Fig. 6F). However, sporulation was greatly reduced by inducing N-deg-Rim4-V5 removal between 14.5h and 17.5h in SPM (Fig. 6B). This finding emphasizes the physiological importance of programmed Rim4 degradation by autophagy, and led us to investigate the fate of the mRNAs that were released prematurely due to induced N-deg-Rim4-V5 removal.

During meiosis, the level of intracellular *CLB3* mRNAs reaches a peak at ~15h (in SPM) and remains stable in the next ~4 hours (Fig. 6E); therefore, we induced degradation of N-deg-Rim4-V5 at 15h (in SPM) and measured the *CLB3* at 17h (in SPM) by RT-qPCR, reasoning that 2h is sufficient for Rim4 degradation (Fig. S6A). Remarkably, using *ACT1, POR1, and PGK1* as internal control, we found that induced N-Deg-Rim4 degradation reduced the intracellular level of *CLB3* mRNAs in an autophagy-dependent manner (Fig. 6G). While being more abundant than *CLB3* (Fig. S6C), unlike *CLB3*, the *ACT1, POR1, and PGK1* are not targets of Rim4-mediated translational suppression. Consistently, the protein levels of Pgk1 and Por1 are relatively constant during meiosis (Fig. S6D; S6E). Therefore, our finding indicates that Rim4 specifically protects its target mRNAs from autophagy. As further support that mRNAs released from Rim4 are sensitive to autophagy, we found that *CLB3* mRNAs in *Δrny1* cells after induced N-deg-Rim4 degradation remains stable and shows no synthetic effect with 1-NM-PP1 inhibited autophagy (Fig. 6H). Rny1 is involved in vacuolar RNA degradation^23,45^; therefore, this result demonstrates that Rim4 protects bound *CLB3* mRNAs from autophagy-mediated vacuolar delivery.

Lastly, we examined the stability of a list of Rim4-sequestered mRNAs clustered with *CLB3* based on previous studies^18,46^. Most *CLB3*-clustered mRNAs showed reduced intracellular levels 2 to 3 hours after induced N-deg-Rim4 degradation (Fig. S6F). Interestingly, Rim4 also stabilizes RIM4 mRNAs against autophagy (Fig. S6F), indicating that the programmed rapid removal of Rim4 is accompanied by reduced Rim4 biogenesis due to *RIM4* degradation. Therefore, we conclude that the strictly programmed Rim4 degradation by autophagy prevents unwanted degradation of Rim4-sequestered mRNAs by autophagy.

In summary, we demonstrate that Rim4 binds mRNAs in the nucleus as a prerequisite for nuclear export. The Rim4-mRNA complex is resistant to Atg11-dependent selective autophagy until the programmed mRNA dissociation from Rim4 during meiotic divisions. Upon dissociation, both Rim4 and the mRNAs are sensitive to autophagy. Thus, we show evidence that autophagy inspects the timing of Rim4-mRNA complex assembly and disassembly, which is essential for meiosis and sporulation.

## Discussion

### The role of Rim4 in nucleus

Rim4 is essential for meiosis and sporulation in budding yeast. Rim4 expression is controlled by *IME1*, an early meiosis transcription factor^18,24^. Since its expression, Rim4 accumulates as a suppressor of translation in the cytoplasm until its rapid removal by autophagy and proteasome at the end of meiosis I^6,16,19^. However, the subcellular distribution of Rim4 and the associated functional relevance remains elusive. This study found that Rim4 dynamically shuttles between the nucleus and cytosol with an NLS that facilitates its nuclear import. So, why Rim4 enters the nucleus, the compartment of transcription? Our data indicate that Rim4 binds mRNAs in the nucleus as a prerequisite of its nuclear export; Rim4 variants that failed to bind mRNAs, e.g., Rim4(FLm), yield nuclear retention phenotype (Fig. 2). Among the family of RBPs, this phenotype is not unique to Rim4. Forming a complex with RNAs is required for efficient nuclear export of many RBPs that primarily reside in the cytoplasm^47,48^, including Pab1, which binds to the common moiety of all eukaryotic mRNAs, i.e., poly(A) tail, and controls mRNA deadenylation and translation initiation^26–28^. Thus, entering the nucleus allows the master RBPs, e.g., Rim4 and Pab1, to control a broad range of mRNA metabolism at the earliest possible stage. In addition, the genetic deletion of Rim4 abolishes meiotic DNA replication, yet, the mechanism is unknown. Now we know that Rim4 is present in the nucleus; therefore, Rim4 may take part in RNA biology in the nucleus, independent of its role as a cytosolic translation suppressor.

Due to nutrient depletion, yeast meiosis features active autophagy. One benefit of the nucleocytoplasmic shuttling of Rim4 is that this protects Rim4 and Rim4-sequestered mRNAs from autophagy. Specifically, Rim4 and mRNAs in the nucleus are protected from autophagy by the nuclear membrane; Rim4 relocation to cytoplasm requires the Rim4-mRNA complex assembly, which ensures its resistance to autophagy. Given the importance of Rim4 nucleocytoplasmic shuttling, our accompanying manuscript has elucidated a phosphorylation-based mechanism that controls Rim4-mRNA interaction, subcellular distribution, and function. Future studies should clarify the spatial Rim4 interactions (nucleus versus cytoplasm) and whether other RBPs, e.g., Pab1, adopt a similar mechanism to survive active autophagy in the cytosol.

### The mechanism of autophagic Rim4 degradation

Autophagy typically uses receptors to target its substrates, e.g., aberrant forms of proteins and damaged or unwanted organelles^38,49,50^, which can activate Atg1 kinase in concert with the biogenesis of autophagosome that eventually engulfs selected cargoes^41^. More than 30 autophagy receptors have been identified^51^; the list is growing. Our data demonstrate that the selective Rim4 degradation by autophagy relies on Atg11. Atg11 is a scaffolding protein that serves as a hub or platform for recruiting the core autophagic machinery to initiate selective autophagosome formation^52^. Besides, the interaction of Atg11 with a selective autophagy receptor (SAR) associated with a specific substrate, e.g., Atg32 as a SAR for mitophagy^53,54^, recruits the selected substrates to the PAS. Thus, to become a selective autophagy substrate, Rim4 might bind an unknown SAR; alternatively, Rim4 could interact with Atg11 directly. Torggler et al. revealed that tethering Atg11 to the cargo is sufficient to recruit the cargo to the PAS, bypassing SARs^55^. Notably, in most known SARs, the region that binds to Atg11 requires phosphorylation of certain serine(s) prior to Atg11 binding. The cumulative phosphorylation has been found to stimulate proteasome-mediated Rim4 degradation^19^. In our accompanying manuscript, we have defined several regions of Rim4 in which phosphorylation and dephosphorylation showed distinct meiosis and sporulation phenotypes. Next, it will be crucial to examine whether and how phosphorylation at specific phosphorylation sites (P-sites) of Rim4 allows the temporal degradation of Rim4, but not Rim4-mRNA, via the Atg11-dependent selective autophagy.

Rim4 contains three conserved RRMs spaced by low complexity sequences (LCD), including two poly(N) prion-like regions that mediate Rim4 self-assembly^16,18^. Recombinant Rim4 stimulates Atg1 kinase activity *in vitro*, indicating that the intrinsic property of Rim4 is sensitive to autophagic degradation. Many RBPs contain RNA recognition motifs (RRM). RRM is one of the most common eukaryotic protein motifs, encoded in ~2% of all human genes^32^. We found that the RRMs of Rim4 are likely involved in activating Atg1: recombinant Rim4(ΔC) that lacks LCD, and the RRM-containing Pab1 activates Atg1 *in vitro* (Fig. 5D; 5E). Due to the conservation and popularity of the RRM sequence, it is vital to examine whether the consensus sequence/structure of Rim4 RRM provides a principle for autophagy-mediated degradation of other RRM-containing RNA binding proteins.

Despite the role of RRMs, the possible involvement of LCD sequence in autophagic Rim4 degradation needs to be evaluated in the future. The poly(N)-containing LCD sequence is required for high-order Rim4 self-assembly and is under phosphor-regulation^16,19^. Therefore, we suspect that a controlled size of Rim4 self-assembly might facilitate RRM-stimulated Atg1 activation by recruiting a set of locally concentrated Atg components to initiate PAS assembly. In the simplest scenario, since dimerization or oligomerization of Atg1 is a mechanism for its activation^56,57^, accumulation of Atg1 on the self-assembled Rim4 alone might promote Atg1 clustering and Atg1 activation. In line with this model, the mRNA release might be required for Rim4 self-assembly into the right size, which optimizes its chance of being recognized by autophagy.

### The physiological relevance of autophagic Rim4 degradation

Rim4 is degraded by both autophagy and proteasome^6,19^; nonetheless, autophagy is essential and irreplaceable for timely translation of Rim4-sequestered mRNAs, e.g., CLB3, even in the presence of proteasome activity^6^. On the other hand, due to its general inaccessibility to autophagy, nuclear Rim4 degradation might solely rely on the proteasome. Interestingly, the Rim4(47E) that mimics the cumulative phosphorylation of the whole LCD stimulates proteasome-mediated Rim4 degradation^19^. In our accompanying manuscript, we used a mutagenesis approach showing that Rim4 nuclear localization increased as a response to cumulative phosphorylation to several specific regions of the Rim4 LCD domain; in addition, we showed that Bmh1/2, the yeast 14-3-3 proteins, bind and protect Rim4 from autophagy in a site-specific phosphorylation-dependent manner, while Bmh1/2 binding was previously reported to stimulate Rim4 degradation by the proteasome^58^. These findings suggest that the two conserved eukaryotic degradation pathways might target different forms of Rim4 or in different subcellular compartments. Thus, one future direction is to investigate this possibility in the context of meiotic translation.

One significant finding in this study is that autophagy spares the Rim4-mRNA complex. Thus, Rim4 shelters its sequestered meiotic mRNAs to secure their scheduled translation at a later stage. Historically, the autophagy-RNA interface was understudied. Previous studies documented that autophagy degrades RNAs in various organisms^20,22,23^, yet, we know little about related regulation or the physiological relevance of such activity. Recently, MaKino et al. reported that autophagy triggered by TORC inhibition (Rapamycin treatment) prefers ribosome-associated mRNAs^23^, while it is unclear how autophagy picks these mRNAs or spares the others. Interestingly, our study finds that induced Rim4 (N-Deg-Rim4) removal destabilized *CLB3* mRNA and mRNAs clustered with *CLB3* (targets of Rim4) while activating the translation. Thus, in the simplest scenario, Rim4 protects its RNA substrates from autophagy by sequestering them away from active ribosomes. This might explain why the timing of Rim4-mRNA complex assembly and disassembly is essential for meiosis and sporulation. Nonetheless, the mRNA consensus of Rim4 target is unknown, the range of Rim4-controlled mRNA stability on meiotic translation, and hence meiosis, is yet to be defined.

What is the functional orthologue of Rim4 in higher eukaryotic systems? A previous study suggested that DAZL, a germ-cell-specific RBP essential for gametogenesis, behaves similarly to Rim4, i.e., forming amyloid-like aggregates in mice gametogenesis. Here, we found that Rim4 is like DAZL in terms of binding to poly(A) binding protein (Pab1)^59^. Pab1 is a conserved master mRNA (poly [A]) binding protein that stabilizes mRNAs and promotes their translation^28,60^. Notably, Pab1 is the first RBP characterized in complexes with Rim4. However, it is unknown how Rim4-Pab1 interaction affects meiotic RNA biology.

## Supplemental Figures

**Figure S1. NLS-containing Rim4 resides in the nucleus and the cytoplasm**

A) Schematic of induced expression of *NDT80* (*pGAL:NDT80*, for synchronized metaphase I entry) by 1μM β-estradiol. 1μM β-estradiol was added to meiotic cells after 12 h in SPM to synchronize the entry of meiotic divisions. Synchronized cells entered metaphase I, anaphase I, metaphase II, and anaphase II sequentially.

B) The representative FM images of prophase I meiotic cells expressing EGFP-Rim4. Before imaging, the meiotic cells were stained with cell membrane-permeable RedDot™2 Far-Red Nuclear Staining dye (diluted from 200X stock) for 10 minutes at room temperature (RT). The white dashed circle marks the nucleus. The scale bar is 5 μm.

C) The LocTree3 prediction^29^ of Rim4 C terminus subcellular localization. Support Vector Machine method was used to predict the subcellular localization: 1) membrane (0.07) or nonmembrane (0.93), 2) secretory pathway (0.05) or non-secretory pathway (0.95), 3) organelle localization (0.01) or non-organelle localization (0.99), and 4) cytosolic (0.05) or nucleic (0.95). Based on the scores (shown above in the brackets), the Rim4 C terminus was predicted to be localized in nuclei.

D) The sporulation efficiency of indicated Rim4 variants, presented as the tetrads percentage; more than 300 cells were counted for each strain. *NDT80* expression was induced at 12h.

E) Top, Schematic illustration of the Rim4 protein sequence. Bottom, A PyMOL cartoon representation of the 3D structure Rim4 (S. *cerevisiae*) predicted by Alpha-fold. The NLS site is framed in a red box with residues highlighted in red. Domains, including RRM1, RRM2, and RRM3, were labeled in orange, and C terminal LCD was labeled in green.

**Figure S2. Rim4 binds mRNAs in the nucleus**

A) Top, the schematic of the Rim4 protein harboring mutated RRM1 and RRM3, depicted. Bottom, A PyMOL cartoon representation of the 3D structure Rim4 predicted by Alpha-fold (full length) or Robetta (RRMs). The mutated RNA binding sites, i.e., F_96_ and F_349_, with residues highlighted (cyan), are viewed in separate zoom-in windows (right).

B and C) the representative FM images of meiotic cells at 12h in SPM, expressing the indicated EGFP-Rim4 variants. The truncation of the C-terminus (ΔC289) or fusing an NES to the N-terminus rescue the nuclear retention of Rim4(FLm) or Rim4(ΔN424). Scale bar, 5 μm.

D) Percent of cells forming spores (tetrads) after 60 hours in SPM. As indicated, the cells harbor one allele of EGFP-Rim4 WT with either one allele of Δ*rim4* or one allele of EGFP-Rim4(Δ*N424*). NDT80 expression was induced at 12h in SPM.

**Figure S3. Rim4 dissociates from mRNAs during meiotic divisions**

A Schematic illustration of *the CLB3* reporter construct (top), CLB3 reporter transcription and translation (bottom left), and the *CLB3*-luciferase assay (bottom right). Top, the *CLB3* reporter contains a Ubiquitinated N-degron to accelerate its turnover by proteasome so that *CLB3*-luciferase expressed during previous mitotic cell cycles can be depleted before entering meiosis.

Bottom left, before meiotic divisions, Rim4 will bind to the CLB3-5’-UTR of the report mRNAs to suppress translation; during the meiotic divisions, the reporter mRNA was released and translated into protein, i.e., nano-luciferase (Ubi-NLuc). Bottom right, the cell lysate containing Ubi-NLuc converts NLuc substrate (furimazine) into furimamide and glow-type luminescence.

B) Whole cell extracts derived from *GAL-NDT80* synchronized meiotic cells harboring EGFP-Rim4 and CLB3-Flag, at indicated time points in SPM were analyzed by immunoblotting (IB) with indicated antibodies. Arrow, *NDT80* induction at 12h in SPM. +AI (Autophagy inhibitor), 5μM 1NM-PP1 was applied at 12h in SPM. Hxk1, loading control.

C and D) FM analysis of synchronized meiotic cells expressing EGFP-Pab1 and mScarlet-Atg8. (C) Shown are representative images; white arrow, mScarlet-Atg8 puncta that do not co-localize with EGFP-Pab1 (n≥300 cells, 3 replicate experiments). Scale bar, 5μm. (D) percentage of cells with Atg8/Pab1 co-localization. Only cells that carry EGFP-Pab1 puncta were analyzed here. (n≥300 cells, 3 replicate experiments, t-test).

**Figure S4. Selective autophagy degrades Rim4 and is important for sporulation**

A and B) IB analysis with indicated antibodies (A) of cell lysate and quantification (B) of signal intensity showing EGFP-Rim4, Ape1-EGFP, and free GFP protein level during sporulation at indicated time points in SPM. (A) Top, the schematic of a cell harboring one allele of EGFP-Rim4 and one allele of Ape1-EGFP under *the RIM4* promoter. Bottom, representative IB images. Pgk1, loading control. NDT80 expression was induced at 12h in SPM. (+) Autophagy, mock treatment;

(-) autophagy, 5μM 1NM-PP1 treatment at 12h in SPM. (B) Graph showing the ratio of EGFP-Rim4/ Ape1-EGFP IB signal. AU, arbitrary units.

C) Percent of cells forming spores (tetrads) after 60 hours in SPM. As indicated, the cells harbor ATG11 (WT) or *Δatg11*. NDT80 expression was induced at 12h in SPM to enable meiotic divisions and sporulation.

**Figure S5. Recombinant proteins in this study(A)** Commassie blue staining image showing the purified recombinant proteins from E.Coli.

B, C and D) FM analysis of synchronized meiotic cells expressing mScarlet-Atg8 and EGFP-Rim4. 5uM 1NM-PP1 was applied at prophase I (12h in SPM) if necessary. A) Representative images: Yellow arrows show the co-localization of mScarlet-Atg8 and EGFP-Rim4; white arrows show the puncta not co-localized. B) percentage of cells with Rim4/Atg8 co-localization. Only cells that carry EGFP-Rim4 puncta were analyzed here, (n≥300 cells, 3 replicate experiments, t-test).

C) percentage of cells with Rim4 puncta in total cells. (n≥300 cells, 3 replicate experiments, t-test).

**Figure S6. Induced Rim4 degradation by proteasome reduced intracellular stability of Rim4-sequestered mRNAs**

A) IB analysis showing cellular levels of Rim4-V5 and N-Deg-Rim4-V5 along meiosis at indicated time in SPM. After 2h of β-estradiol-induced TEV expression, the cellular level of N-Deg-Rim4-V5 robustly decreased. IB, α-V5.

B) The sporulation efficiency in cells expressing Rim4-v5 or N-Deg-Rim4-V5, presented as the tetrads percentage; more than 300 cells were counted for each strain.

C) Graph of *CLB3* mRNA levels (ΔCt analysis) in cells, relative to indicated *mRNAs*. At 17h in SPM, cells were collected for RNA extraction, cDNA reverse-transcription, and qPCR. Note that the ratio of *CLB3/CLB3* reporter is close to 2 because the latter is transcribed from only one allele of the gene.

D and E) IB analysis showing constant cellular levels of Por1 (E) and Pgk1 (F) along meiosis at indicated time in SPM. IB, α-V5.

F) Heat map showing the change of intracellular level of indicated mRNAs (RT-qPCR), responding to N-Deg-Rim4 degradation. 1 μM β-estradiol was added to SPM at 15h to trigger N-Deg-Rim4 degradation. Green colors (<1 and ≥0.091) and red colors (≤4.5 and >1) indicate reduced and enhanced, respectively, mRNA levels in the cells 2h after induced N-Deg-Rim4 removal. The black color indicated no change (=1).

