## Supplementary figures and images for "Autophagy inspects Rim4-mRNA interaction to safeguard programmed meiotic translation"

### Figure S1

Zhang et al, Figure S1

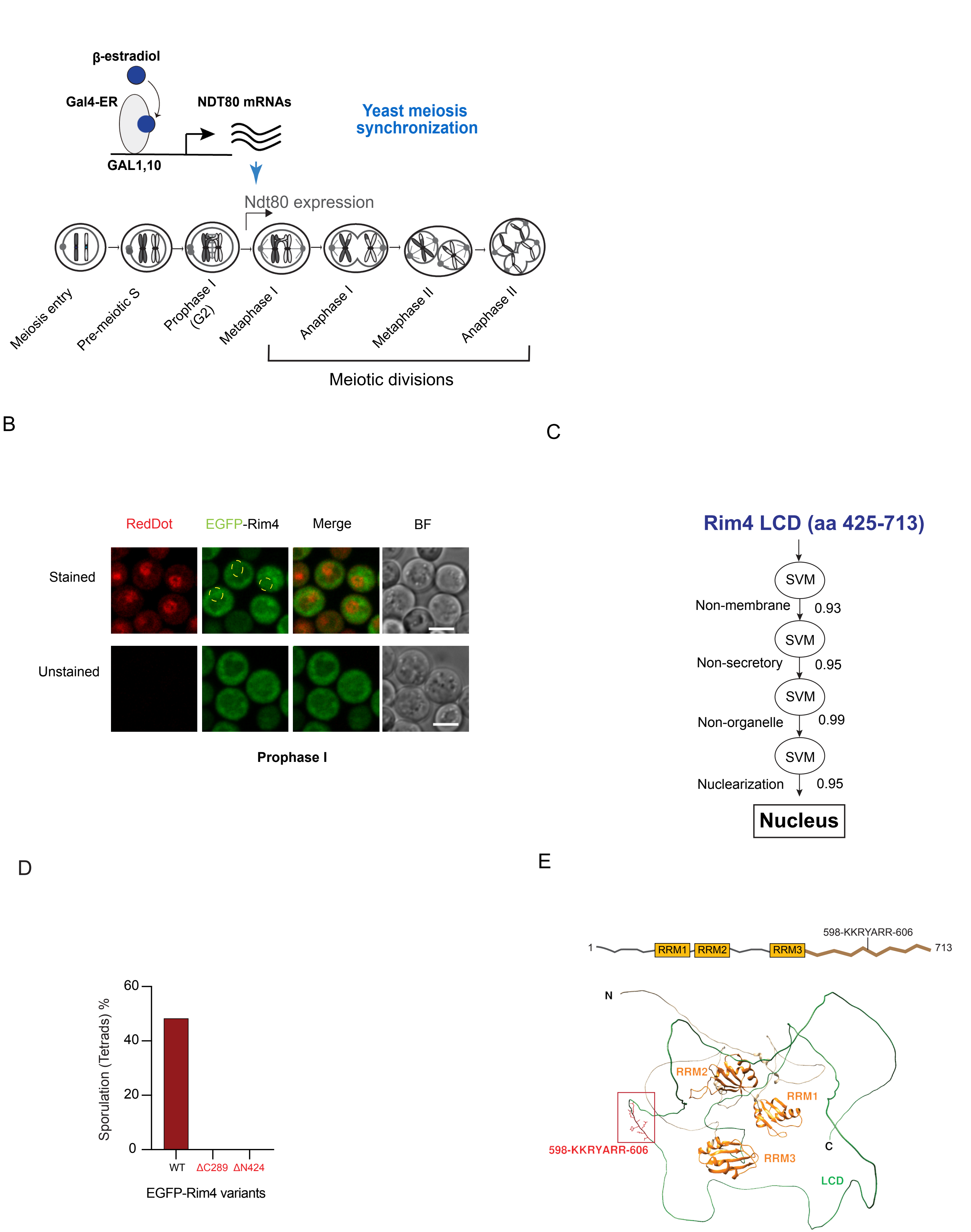

### Figure S2

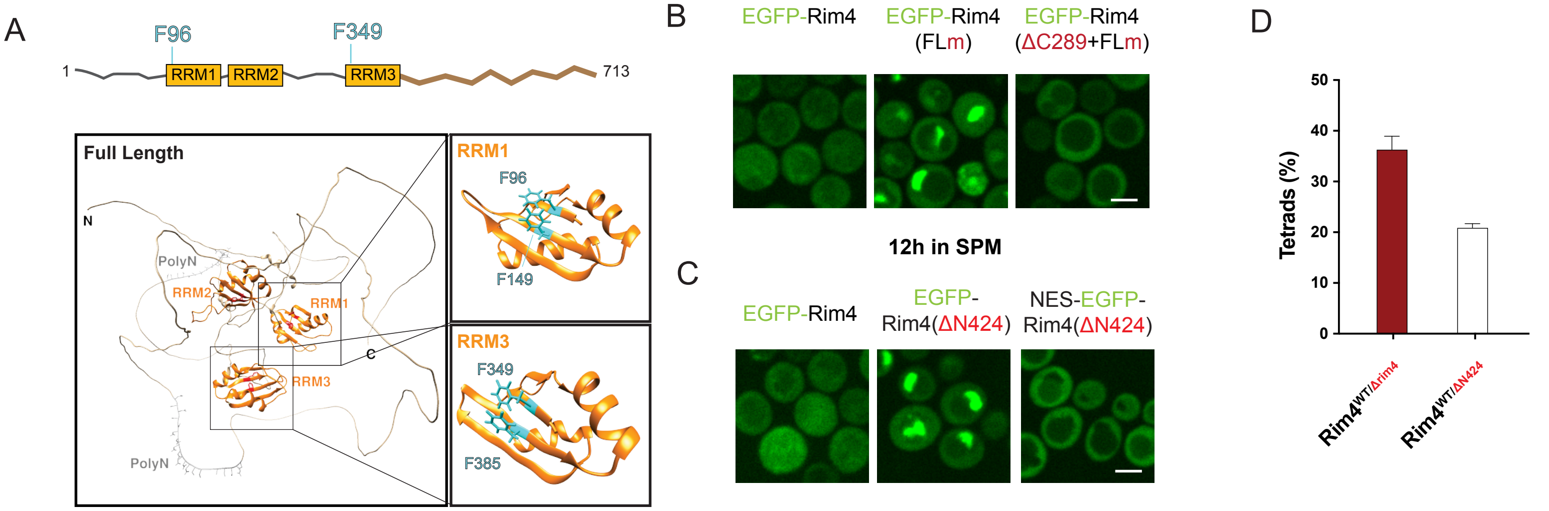

### Figure S3

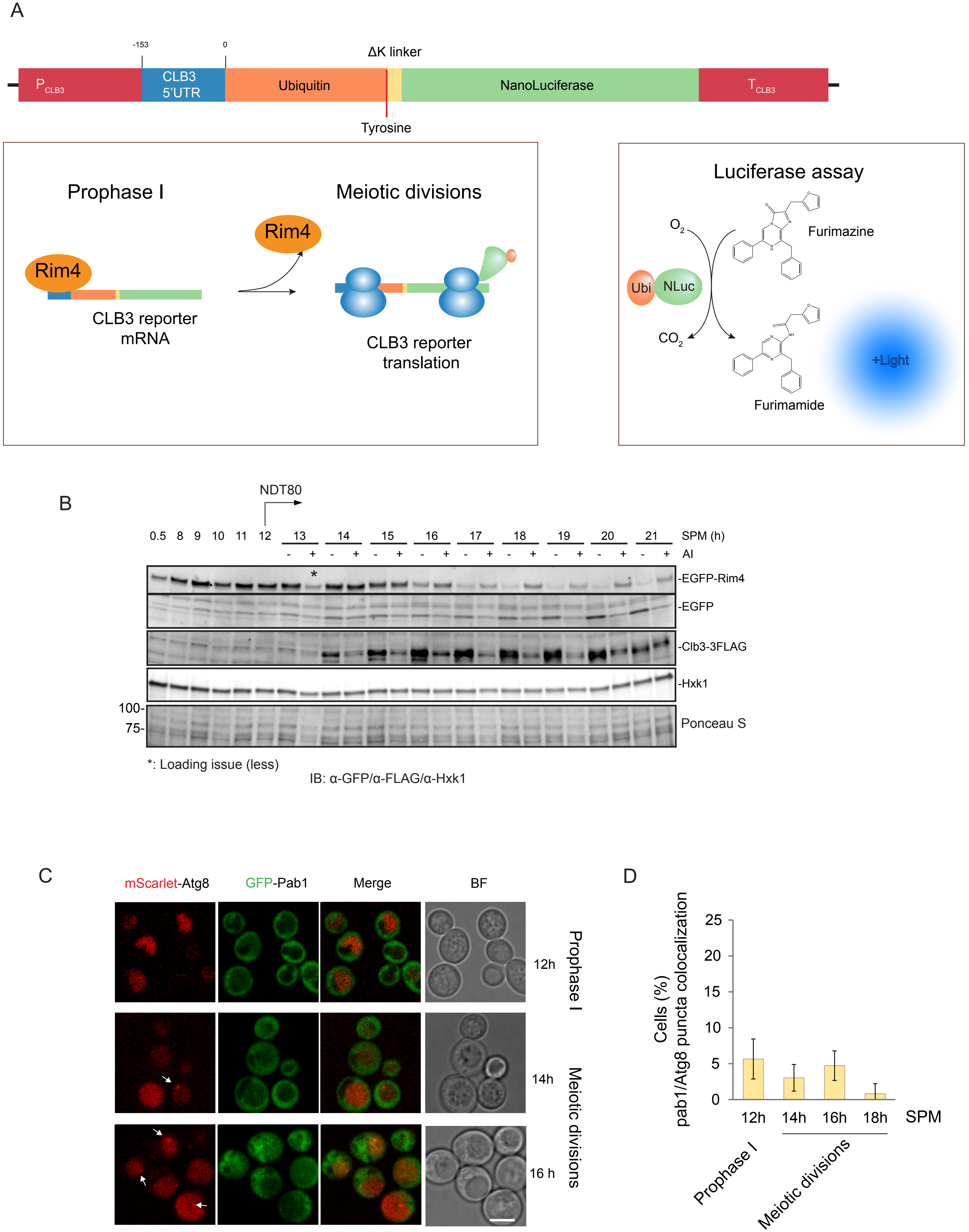

### Figure S4

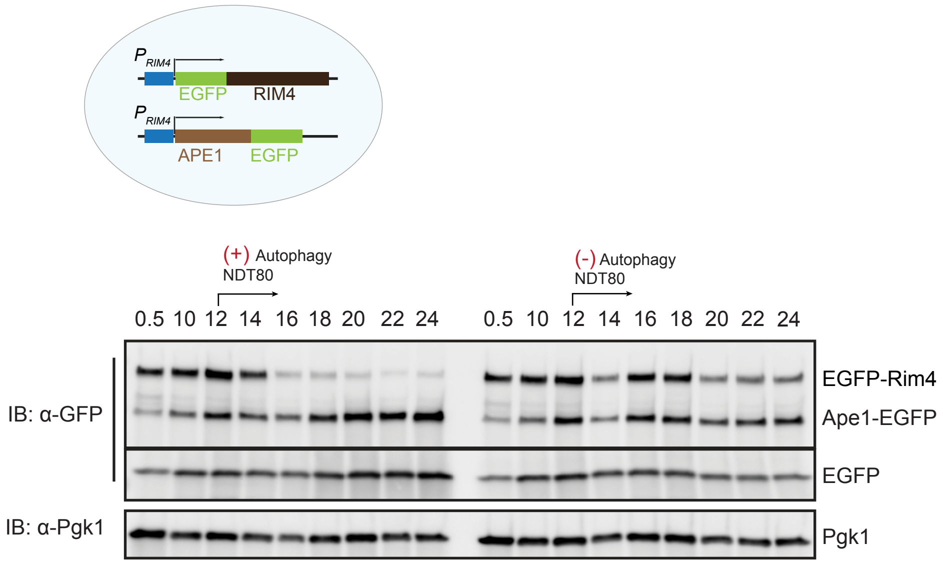

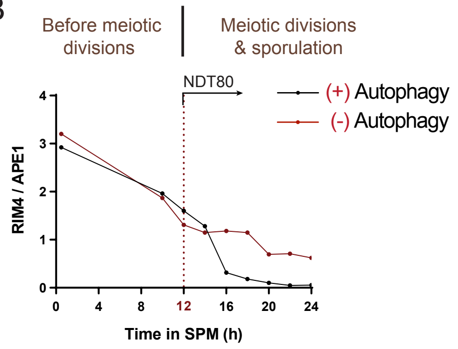

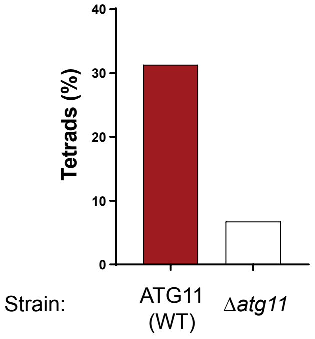

### Figure S5

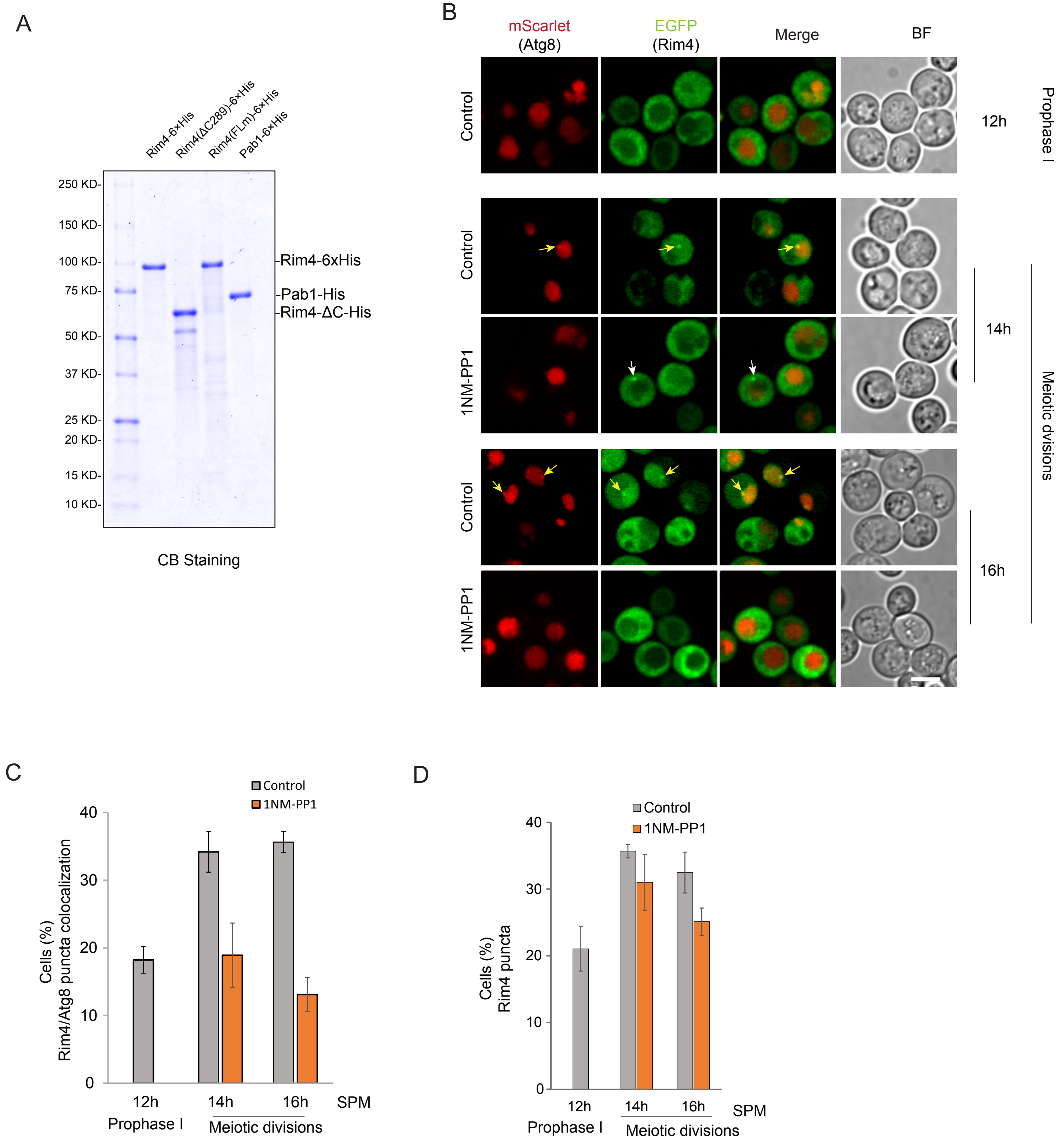

### Figure S6

Zhang et al, Figure S6

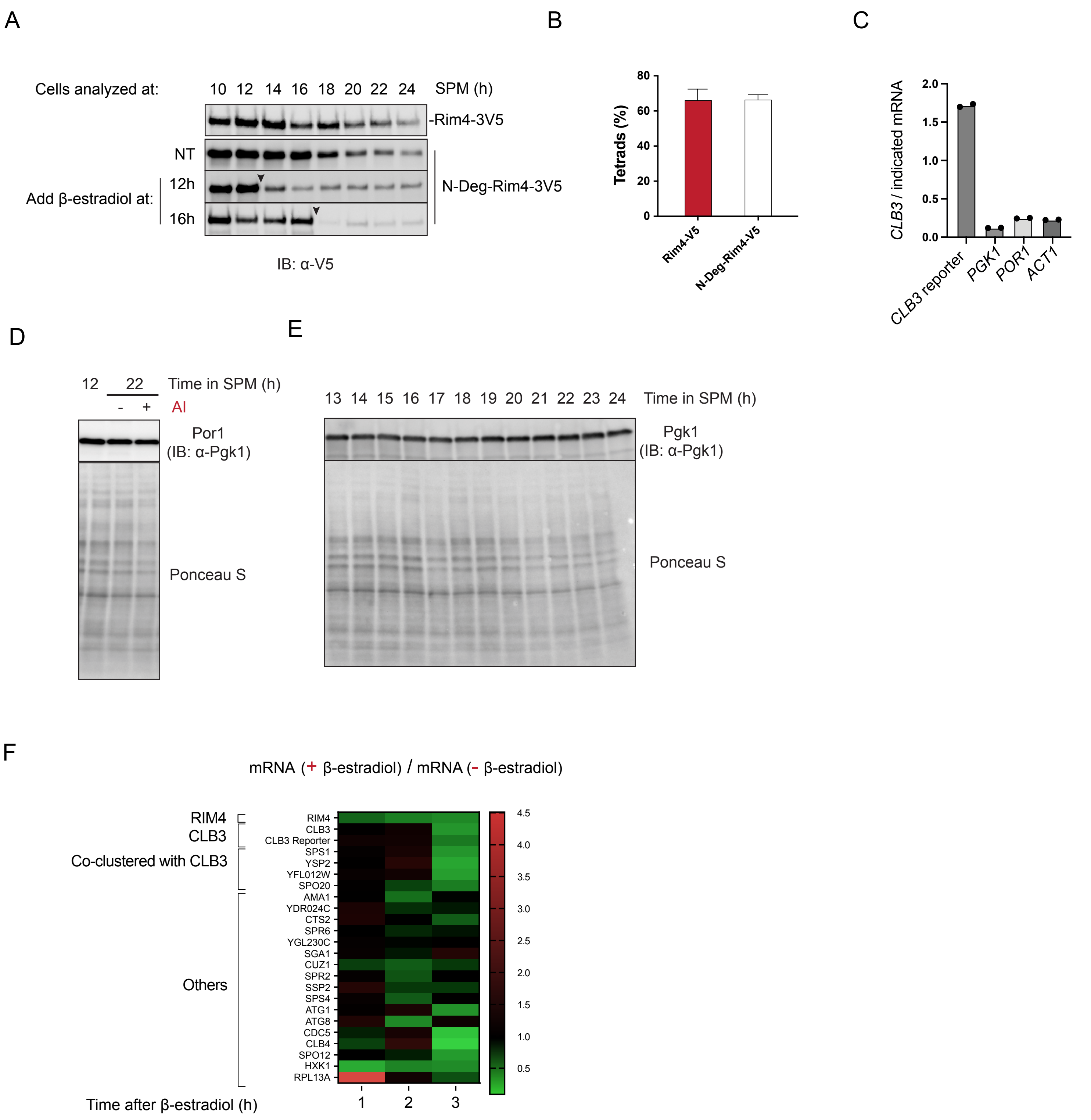
