## Supplementary material for "Autophagy inspects Rim4-mRNA interaction to safeguard programmed meiotic translation": Methods

**Yeast**

Saccharomyces cerevisiae strains are derivatives of W303 (ade2-1 his3-11,15 leu2-3,112 trp1-1 ura3-1 can1-100) (Table S1). Note that, unless otherwise indicated, the analog sensitive version of Atg1(Atg1-as) was created by gatekeeper residue change (M102G) as described earlier (1, 2), introduced into the background of listed stains to allow conditional autophagy inhibition by 1NM-PP1.

Deletion strains were constructed in a parent background by PCR-mediated knock-out with one of the following drug resistance or prototrophic marker cassettes: pFA6a-kanMX6/pFA6a-NAT (Longtine et al., 1998), pKlURA/pCgHIS (Goldstein and McCusker, 1999). C- or N- terminal tagging at the endogenous genomic locus was introduced by PCR-mediated epitope tagging as previously described (Denic and Weissman, 2007). Specifically, strains with GAL-NDT80 GAL4-ER were constructed by replacing the endogenous NDT80 promoter with the inducible GAL1,10 promoter as described (Carlile and Amon, 2008). Strains with ndt80Δ was constructed with pFA6a-NAT marker cassette replacement of NDT80 coding sequence.

The N-Deg-Rim4 strains are created by the following steps: first, the plasmid containing P_ACT1_-Zif268(DBD)-hEsR-VP16 (ZEV) cassette and P_ZEV_(modified GAL1 promoter with 6xZif268 binding sites)- Tobacco Etch Virus protease (TEV) (pFW43) was linearized and integrated into the yeast genome at *leu2* locus and selected by leucine prototrophy. Endogenous Rim4 was replaced by N-Deg-RIM4 through *URA3* cassette mediated knock-out and 5’-FOA resistance selection. The N-Deg-RIM4 was tagged with a 3xV5 tag at C-terminus for epitope detection if necessary. The pCLB3-CLB3 5’-UTR-UbiYΔΚ-NanoLuciferase (CLB3 reporter) carried by a pRS303 backbone plasmid was linearized and integrated into *his3* locus. Plasmids with Nup49-mScarlet, mScarlet-Pab1 or EGFP-Pab1in pRS304, mScarlet-ATG8 in pRS303, driven by their endogenous promoters, were integrated into the genomic locus of *his3/trp1*. Strains harboring EGFP-Rim4 mutants were made by introducing mutated pRS303-harbored EGFP-Rim4 into *his3* locus of parent strains, in which endogenous *RIM4* coding sequence was deleted by PCR-mediated knock-out with *URA3* marker cassette.

**Media**

The following media were used in this study. YPD (2% peptone, 1% yeast extract, 2% glucose), YPA (2% peptone, 1% yeast extract, 2% potassium acetate), SD (0.67% yeast nitrogen base, 2% glucose, auxotrophic amino acids and vitamins), and standard sporulation medium SPM (0.6% potassium acetate, pH=8.5). SD-dropout medium was made with dropout stock powder lacking histidine, tryptophan, leucine, uracil and/or methionine.

**Plasmid construction**

Plasmids used in this study are listed in Table S2.

pRS303-EGFP-Rim4 was constructed by subcloning PCR amplified regions of genomic Rim4 with its promoter and terminator into pRS303. EGFP fragment was fused to the N-terminus of RIM4 by Gibson assembly cloning. Mutagenesis on EGFP-Rim4, as shown in Table S2, was introduced by overlap extension (OE) PCR.  pRS304- Nup49-mScarlet was constructed by subcloning PCR amplified regions of genomic NUP49 with its promoter and terminator into pRS304. mScarlet, fragment was fused to the C-terminus of NUP49 by Gibson assembly cloning. pRS303-mScarlet-Atg8, pRS304-mScarlet-Pab1 and pRS304-GFP-Pab1 were constructed in the same way as pRS304-NUP49-mScarlet except that the mScarlet/GFP fragment was fused to the N-terminus of target gene.

Vectors for bacterial expression of 6His-3FLAG-tagged Cdc14 were created by subcloning PCR amplified regions of the genomic Cdc14 encoding region into the NdeI/XhoI restriction sites of pET28a.

NanoLuciferase, as a CLB3 expression reporter, was driven by CLB3 promoter (133 bp). The Clb3 5’-UTR sequence (153 bp) ^1^ was added before the reporter ORF. An N-Degron fragement, reported in previous researches: UbiYΔK ^2^ was fused to the N terminal of the NanoLuciferase coding region. The NanoLuciferase coding region was amplified by PCR from the plasmid pDONR221-NanoLuc, a gift from Vladimir Denic Lab.

**Sporulation**

A single colony of yeast strains from YPD plate were picked, spread on YPG (3% glycerol) plates and grown at 30°C for 2 days. Colonies grown out were spread to YPD plates and grown until cells formed a lawn (~ 24 hours). Cells on plates were collected and suspended in YPA liquid medium (OD_600_=0.3) and grown for 14-16h at 30°C. Cells were then pelleted, washed with water twice and resuspended in SPM to final OD_600_=2. For M-phase synchronization, following incubation in SPM for 12 hours, strains containing GAL-NDT80 GAL4-ER were released from the prophase I arrest by addition of 1 μM β-estradiol to induce Ndt80 expression. To assess the percent spore formation, cells with matured spores were counted after 48 hours in SPM at room temperature. At least 300 cells for each strain were counted under bright field with Olympus microscope (BX40, 40x objective).

**Meiosis Synchronization by Ndt80 Arrest/Release**

For synchronizing NDT80-in cells, following incubation in SPM for 12 hours, strains containing GAL-NDT80 GAL4-ER were released from the prophase I arrest by addition of 1μM β-estradiol (10mM stock in ethanol, Sigma E2758-1G). To inhibit Atg1-as, 5 μM of 1-NM-PP1, was added (10 mM stock in DMSO) to SPM.

**DNA Replication Analysis with Flowcytometry**

Cells were collected at 12 hr in SPM and fixed in 70% ethanol at 4˚C overnight. Afterward, the cells were pelleted and resuspended in 50 mM Sodium Citrate (pH 7.0), followed by sonication at 30% power for 15 sec. Cells were pelleted again and resuspended in the same solution. The cells were treated with 0.25 mg/mL RNase A at 37˚C overnight. Next, the cells were incubated in 1 µM SYTOX Green dye at RT in the dark for at least 1 hr before being analyzed on a flow cytometer (BD FACSCalibur). The forward scatter (FSC), and side scatter (SSC) were used to gate the living cells. The cell counts of green (530/30 nm) signal in the gate were collected and analyzed by Flowing Software 2.5.1 (<https://bioscience.fi/services/cell-imaging/flowing-software/>).

**Immunoblotting**

1.4 OD_600_ of yeast cells were collected by centrifugation, resuspended in 100µl 2x SDS sample buffer and boiled at 70°C for 5 min [2xSSB: 62.5 mM Tris-HCl, pH=6.8; 2% SDS; 0.05% BPB; 10% Glycerol; 5% 2-Mercaptoethanol; 1× Protease Inhibitor Cocktail (Roche, 11873580001), 1 mM PMSF]. Proteins were separated by SDS-PAGE (90 min at 150V) using 4-20% Criterion™ TGX Stain-Free™ gel (Bio-Rad, 5678095) and electro-blotted onto nitrocellulose membranes (Bio-Rad, 1620115) using a Trans-blot SD semi-dry transfer cell (Bio-Rad, 1703940). After blocking with 5% skim milk or 1% BSA in TBST and incubated with respective primary antibodies followed by HRP-, StarBright® B700- or Alexa Fluor 488-conjugated secondary antibodies. For detection and quantitative analysis, the IB images were captured by ChemiDoc™ MP imaging system (Bio-Rad, 12003154), and analyzed using Image Lab™ (Ver. 6.0.1) software (Bio-Rad).

We used the following antibodies: Recombinant monoclonal anti-Thiophosphate ester rabbit IgG [51-8] (Abcam, ab92570), monoclonal anti-GFP mouse IgG (Roche, 11814460001, 1:5000), polyclonal anti-Hexokinase 1 rabbit IgG (United States Biological, 169073, 1:10000), polyclonal anti-Ndt80 rabbit IgG (1:10000), monoclonal anti-HA mouse IgG1 (Thermo Fisher, 21683, 1:3000), monoclonal anti-FLAG M2 mouse IgG (Millipore-Sigma, F3165, 1:5000), polyclonal anti-FLAG rabbit IgG (Millipore-Sigma, SAB4301135, 1:5000), StarBright® B700 labeled goat anti-mouse IgG secondary antibody (Bio-Rad, 12004158, 1:5000), HRP conjugated goat anti-mouse IgG (Bio-Rad, STAR207P) and StarBright® B700 labeled goat anti-rabbit IgG (Bio-Rad, 12004161, 1:5000). Mouse-anti-puromycin 12D10 monoclonal antibody (Sigma, MABE343, 1:5000).

**Recombinant protein purification**

pET-based protein expression in BL21 (DE3) E. coli cells was induced by IPTG as described previously (Wang et al., 2010). 6His-Rim4/ 6His-Pab1 were expressed at 16°C overnight. Cells were collected by centrifugation, resuspended in bacteria lysate buffer (BLB: 20 mM Tris-HCl pH 8.0, 250 mM NaCl, 20 mM Imidazole, 10% glycerol, 5 mM 2-ME, 1 mM PMSF), and lysed using a High-Pressure Cell Press Homogenizer‎ (Avestin Emulsiflex-C5). The lysate was supplemented with 0.1 mg/ml DNase I, 1 mM MgCl2, and 0.1% Triton-X 100 and cleared by centrifugation at 10,000g for 10 min at 4°C. The supernatant was applied to an NTA-Ni column (Qiagen, 30230), and washed three times with BLB. The 6xHis- proteins were eluted with 20mM Tris-HCl pH 8.0, 250 mM NaCl, 300 mM Imidazole, 10% glycerol and 5 mM 2-ME. The eluted proteins were separated by Superdex 200 Increase 10/300 GL column (GE Healthcare) equilibrated in size exclusive buffer (SEC: 50 mM Tris/HCl pH 7.5, 250 mM NaCl, 10% glycerol, 5mM 2-ME). Finally, purified proteins were concentrated in SEC and frozen in liquid nitrogen.

**immunoprecipitation**

Cells grown in SPM or YPD medium were pelleted at 3,000 × g for 5 min, 4°C, washed with ice-cold distilled water containing 1mM PMSF. The pellets were resuspended in ice-cold yeast lysis buffer (YLB: 50mM HEPES-KOH, pH 6.8, 150 mM KOAc, 2 mM MgCl2, 1 mM CaCl2, 0.2 M sorbitol, 10mM PMSF, 2x protease inhibitor cocktails [Roche, 11873580001]), dropped into liquid nitrogen, ground using a Retsch ball mill (PM100 or Retsch MM400) and stored at -80 °C for immunoprecipitation (IP), as well as for atg1 kinase assay and immunoblotting (IB).

To isolate the FLAG-Atg1-as complex or Pab1-FLAG, frozen cell lysate powder (~ 500 OD600 units) was thawed in 2ml 1 × IP buffer (50 mM HEPES-KOH (pH 6.8), 150 mM K2OAc, 2 mM MgOAc, 1 mM CaCl2, 15% glycerol, 1% NP-40, 1x protease inhibitor cocktail, 1x phosphatase inhibitors) and cleared twice by centrifugation at 1,000 × g for 5 min at 4 °C. 10 µl of Protein G Dynabeads (Invitrogen) that were loaded with mouse anti-FLAG M2 antibody (Sigma, F3165) were then added to the cell extract, incubated for 3 h at 4 °C with constant agitation. The beads were collected, washed 5 times with IP buffer containing 1% NP-40 and 1 × phosphatase inhibitors, and bound proteins were eluted with 25 µl 1 mg/ml 3 × FLAG peptide (Sigma, F4799) in IP buffer containing 1% NP-40 and 1 × protease inhibitors at 4 °C. Eluates were aliquoted, frozen in liquid nitrogen, and stored at -80 °C. The purified FLAG-Atg1 complex was resolved by SDS-PAGE and analyzed by Sypro Ruby staining (Thermo Fisher, S12000). The purified Pab1-FLAG was resolved by SDS-PAGE and analyzed by FLAG and V5 antibody IB.

**Atg1-as kinase assay**

The cell lysate was mixed with 1 × kinase buffer (150 mM KOAc, 10 mM MgOAc, 0.5 mM EGTA, 5 mM NaCl, 20 mM HEPES-KOH [pH 7.3], 5% glycerol) in equal volume (wt/vol) on ice and thawed by pipetting. After spin at 1000g for 5min twice, supernatants were mixed with different concentration of recombinant Rim4 and equal volumes of 2 × kinase mix (kinase buffer, energy mix [90 mM creatine phosphate, 2.2 mM ATP, 0.45 mg/ml creatine kinase] and 0.2 mM N6-phenylethyl-ATPγS [N6-PhEt-ATPγS, BLG-P026-05]) and incubated for 1.5 h at room temperature. Reactions were quenched with 20mM EDTA and then alkylated with 2.5 mM paranitrobenzyl mesylate (PNBM, Abcam) for 45 min at room temperature, heated in sample loading buffer, and analyzed by SDS-PAGE and following immunoblotting assay. Thiophosphorylated substrates were identified by immunoblotting with a rabbit anti-thiophosphate ester primary antibody [51-8] (Abcam, ab92570) and StarBright® B700 labeled goat anti-Rabbit IgG secondary antibody. Blot imaging was done using a Bio-Rad Chemidoc Imaging System.

12.5ug/ml , 25ug/ml and 50ug/ml recombinant Rim4-His was incubated with 2ul Flag-Atg1-as complex, supplied with 1 × kinase buffer to 15ul, for 20 min in RT, followed by atg1-as kinase assay as described above.

**Live cell nuclear staining**

RedDot™1 Far-Red Nuclear dye (200x, Biotium, #40060) was applied directly to SPM to stain nuclear in live meiotic cells for 5 minutes before capturing images.

**Live cell imaging**

10 µl of cells growing in SD medium or SPM were spread on a slide and imaged immediately with 3i spinning disk confocal microscope, using a 63x oil immersion objective with mRuby filter (561-617/73) and GFP filter (413-525/80). Z-stack (5-10 planes, 0.5 µm/plane) images were acquired if needed. Images were processed using ImageJ. Two-sided t-test was used in the colocalization statistics.

**Yeast total RNA extraction and RT-qPCR**

2 OD yeast cells were collected. The pellets were washed in DEPC water twice and snap frozen in liquid nitrogen. After all samples collected, total RNAs were extracted from the yeast cells with the MasterPure™ Yeast RNA Purification Kit (Biosearch Technologies, MPY03100). Briefly, the cells were resuspended in 300 µL Extraction Reagent for RNA contains 0.2 µg/µL Proteinase K, Incubated at 70°C for 15 min with occasionally vortexing. After cooling down on ice, 200 µL MPC Protein Precipitation Reagent was added to the reaction. Precipitated proteins and cell debris were brought down at 14,000 g for 10 min, 4°C. The supernatant were mixed with 500 µL ice cold isopropanol. The RNA samples were precipitated at -80°C for 30 min, and -20°C for another 30 min. The crude extract of RNAs were spun down at 14,000 g for 10 min, 4°C. Then resuspended the pellets in 200 µL 0.025 U/µL RNase-free DNase I in DNase I buffer at 37°C for 30 min. Sequentially added 200 µL T & C Lysis Solution and 200 µL MPC Protein Precipitation Reagent. Sat on ice for 5 min, and pelleted down at 14,000 g for 10 min, 4°C. The RNAs in the supernatant were precipitated by mixing with 500 µL ice-cold isopropanol, and incubated at -80°C for 30 min, followed by -20°C for 30 min. Brought down the RNA samples at 14,000 g for 10 min, 4°C. The pellets (RNA) were washed with ice-cold 70% ethanol twice. Then air dried the RNA samples and resuspended in 35 µL DEPC water. The concentrations of the RNAs were determined by Nanodrop 2000C.

The RNAs were immediately reverse-transcribed to cDNA after the purification, with the SuperScript™ III First-Strand Synthesis SuperMix Kit (Thermo-Fisher, 18080400). Briefly, 6 µL RNAs was mixed with 0.5 µL Random Hexamer oligonucleotides, 0.5 µL Oligo(dT)_20_ oligonucleotides and 1 µL Annealing Buffer. Incubated at 65°C for 5 min. Cool down for 2 min. Then supplied with 10 µL 2X First-Strand Reaction Mix and 2 µL SuperScript III/RNaseOUT Enzyme Mix. Incubated at 25°C for 10 min, 50°C for 2 hr, 55°C for 30 min and 85°C for 5 min.

The Q-PCR reactions were assembled by mixing 7.5 µL 2x iTaq™ Universal SYBR® Green Supermix (Bio-Rad, 1725121), 3.5 µL Nuclease-free water, 3 µL pre-mixed primer pair (0.25 µM each) (see Table S3) and 1 µL cDNA. The 15 µL samples were loaded to the 96-well reaction plate (Bio-Rad, HSP9655). Ran the samples on a CFX96™ Real-Time PCR Detection System (Bio-Rad, 3600037). The results were analyzed by ΔCt method and normalized to ACT1, unless otherwise indicated, to present the target mRNA level, and ΔΔCt method to compare the same target mRNA level change in two different conditions (e.g. Rim4 removed vs. Rim4 non-treated).

***In vitro* transcription and purification**

The RNA transcription of the 120 nt of CLB3 5’-UTR were synthesized by MEGAscript™ T7 Transcription Kit (Thermo-Fisher, AM1333) and purified by MEGAclear™ Transcription Clean-Up Kit (Thermo-Fisher, AM1908). Briefly, the DNA template containing T7 promoter and the 120 bp of CLB3 5’-UTR DNA were amplified by PCR with a primer pair including the 5’ terminal T7 promoter. Assemble the reaction by mix 1.5 µL DNA template; 2 µL of dATP, dGTP, dCTP and dGTP, respectively; 2 µL 10x Reaction Buffer and 2 µL T7 RNA Polymerase. Incubated the reaction at 37°C for overnight (~14 hr). After the reaction, added 1 µL TURBO DNase to the reaction, and incubated at 37°C for 15 min. Next, brought the transcription reaction volume to 100 µL. sequentially added 350 µL Binding Solution Concentrate and 250 µL 100% ethanol. Transfered the sample to the filter cartridge inserted in the collection tube. Spun down at 14,000 g for 1 min. Washed twice with 500 µL Wash Solution. Eluted with 50 µL 95°C nuclease-free water.

**Electrophoresis mobility shift assay (EMSA)**

EMSA assay was conducted by using the vertical thin (1.5 mm) 1% TAE-agarose gel casted with the Bio-Rad Mini-PROTEAN® casting system (Bio-Rad, 1658022); and sequentially stained with EtBr (to show the nucleic acid) and Coomassie Bright Blue R-250 (to show the protein).

The oligonucleotides of the 120 nt CLB3 5’-UTR were synthesized by Sigma-Aldrich (VC00021), and dissolved to 100 µM with nuclease-free water; the RNA of the 120 nt CLB3 5’-UTR were in vitro transcribed as described above.

5 µM mRNA or 0.5 µM oligoDNA were incubated on ice for 15 min with or without 5 µM recombinant proteins (Rim4-6His or Rim4[FLm]-6His) or same volume of SEC buffer as control. The 5 µM proteins alone were also incubated at the same conditions without the nucleic acids. Then, 1.25 µL EMSA Loading Buffer (200 mM Tris-HCl, 150 mM KCl, 5 mM MgCl2, 0.05% NP-40, 5 mM DTT, 30% Glycerol, 0.3% Orange G) was added to each reaction. Samples were loaded to the 1.5 mm 1% agarose gel in Mini-PROTEAN® vertical electrophoresis cell (Bio-Rad, 1658004) and ran until the yellow dye (Orange G) reached to about 1/3 to the bottom. The gel was first stained in 0.5 µg/mL EtBr solution at RT for 15 min. After washed by flowing water, the gel image was captured by ChemiDoc MP imaging system. Then the gel was stained in Coomassie Bright Blue R-250 solution at RT for 30 min, and de-stained in the de-staining buffer for overnight. The gel image was captured by ChemiDoc.

**Luciferase assay**

Cell lysate preparing: 4 OD_600_ cells harboring the CLB3 reporter were collected in screw-capped 2 mL centrifuge tube. The cell pellets were treated with 10 mM PMSF at RT for 5 min. Then ~50 µL 0.5 mm glass beads and 200 µL Lysis Buffer (100 mM Tris-HCl, pH 8.0, 20 mM NaCl, 2 mM MgCl2, 1% Triton-X 100, 2x cOmplete® Protease Inhibitor Cocktail, 50 µM 2-ME, 2 µg/mL PMSF, 2 mM PMSF) were added to the cell pellets. Smashed the cells by beads-beating for 3x 1 min in cold room, with 1 min rest on ice in each interval. Reverseed the tubes and tapped away the materials form the bottom. Then, the tubes were poked with a 21 G needle at the slope side of the bottom. Inserted each tube into one 15 mL conical tube with one wide tip 1 mL tip inside. Spun down briefly to drain the cell lysates from the 2 mL tubes into the 15 mL conical tubes. Took out the flow through samples with the 1 mL tips in the 15 mL conical tubes to new 1.5 mL tubes. Spin down at 3000 rpm for 5 min to remove the large size debris. The supernatant was used for Luciferase assay immediately.

Luciferase assay: The Luciferase assays were conducted with the Nano-Glo® Luciferase Assay System (Promega, N1110). Briefly, the Nano-Glo® Luciferase Assay Buffer was thawed and equilibrated to RT. Add 1/50 volumes of Nano-Glo® Luciferase Assay Substrate to create the Luciferase Assay Reagent (LAR). Applied 10 µL LAR to 10 µL cell lysate or 10 µL Lysis Buffer as background control in the 96-well white flat bottom reaction plate (Sigma, CLS3917). Mixed by shaking at 567 cpm for 5 s before incubate at RT for 5 min, and then read the results with Synergy H1 plate reader at the reading speed of 100 ms per well. Substated the readings by the background reading(s). And the adjusted values were plotted to graphs.

**Translation monitering by Puromycin labeling (SUnSET assay)**

The assay was described in previous research ^3^. The cells were grown to the time of interest, and puromycin was added to the culture at 10 µg/mL. 2 hr later, the cells were collected for immunoblotting. And new translated proteins, which labeled by Puromycin, were detected by anti-Puromycin 12D10 monoclonal antibody.

1.	Berchowitz, L.E., Gajadhar, A.S., van Werven, F.J., De Rosa, A.A., Samoylova, M.L., Brar, G.A., Xu, Y., Xiao, C., Futcher, B., Weissman, J.S., et al. (2013). A developmentally regulated translational control pathway establishes the meiotic chromosome segregation pattern. Genes Dev *27*, 2147-2163. 10.1101/gad.224253.113.

2.	Houser, J.R., Ford, E., Chatterjea, S.M., Maleri, S., Elston, T.C., and Errede, B. (2012). An improved short-lived fluorescent protein transcriptional reporter for Saccharomyces cerevisiae. Yeast *29*, 519-530. 10.1002/yea.2932.

3.	Schmidt, E.K., Clavarino, G., Ceppi, M., and Pierre, P. (2009). SUnSET, a nonradioactive method to monitor protein synthesis. Nat Methods *6*, 275-277. 10.1038/nmeth.1314.
