## Supplementary material for "Autophagy inspects Rim4-mRNA interaction to safeguard programmed meiotic translation": Table S1

**Table S1**. Strains used in this study. Strains are derivatives of W303 (*ade2-1 his3-11,15 leu2-3,112 trp1-1 ura3-1 can1-100*)

| Strain name | Figures | Genotype | Source |
| --- | --- | --- | --- |
| FWY2331 | 1A, 1E, 1G, S1D, 2B, 2C, 2D, 3J, 3K, 3L | MAT A/α, Atg1(M_102_G)/Atg1(M_102_G), Ndt80::P_GAL1_-Ndt80:KanMX/Ndt80::P_GAL1_-Ndt80:KanMX, leu2::P_ACT1_-Gal4(848).ER:NatMX/leu2::P_ACT1_-Gal4(848).ER:NatMX, ∆rim4::Ura3/∆rim4::Ura3, his3::P_RIM4_-EGFP-Rim4:His3/his3, trp1::P_NUP49_-Nup49-mScarlet:Trp1/trp1 | This study |
| FWY2555 | 1A, 3C, 3D, 3E | MAT A/α, Atg1(M_102_G)/Atg1(M_102_G), Ndt80::P_GAL1_-Ndt80:KanMX/Ndt80::P_GAL1_-Ndt80:KanMX, ura3::P_TDH3_-Gal4(848).ER:Ura3/ura3::P_TDH3_-Gal4(848).ER:Ura3 Rim4::P_RIM4_-EGFP-Rim4:Rim4/Rim4, His3::P_PAB1_-mScarlet-Pab1:His3/ His3::P_PAB1_-mScarlet-Pab1:His3 | This study |
| FWY2556 | 1C, 3I | MAT A/α, Atg1(M_102_G)/Atg1(M_102_G), Ndt80::P_GAL1_-Ndt80:KanMX/Ndt80::P_GAL1_-Ndt80:KanMX, ura3::P_TDH3_-Gal4(848).ER:Ura3/ura3::P_TDH3_-Gal4(848).ER:Ura3, Pab1-3×FLAG:NAT/Pab1, Rim4-3×V5:HIS3/Rim4 | This study |
| FWY2332 | 1D, 1E, S1D | MAT A/α, Atg1(M_102_G)/Atg1(M_102_G), Ndt80::P_GAL1_-Ndt80:KanMX/Ndt80::P_GAL1_-Ndt80:KanMX, leu2::P_ACT1_-Gal4(848).ER:NatMX/leu2::P_ACT1_-Gal4(848).ER:NatMX, ∆rim4::Ura3/∆rim4::Ura3, his3::P_RIM4_-EGFP-Rim4(∆N424):His3/his3, trp1::P_NUP49_-Nup49-mScarlet:Trp1/trp1 | This study |
| FWY2333 | 1D, 1E, S1D | MAT A/α, Atg1(M_102_G)/Atg1(M_102_G), Ndt80::P_GAL1_-Ndt80:KanMX/Ndt80::P_GAL1_-Ndt80:KanMX, leu2::P_ACT1_-Gal4(848).ER:NatMX/leu2::P_ACT1_-Gal4(848).ER:NatMX, ∆rim4::Ura3/∆rim4::Ura3, his3::P_RIM4_-EGFP-Rim4(∆C289):His3/his3, trp1::P_NUP49_-Nup49-mScarlet:Trp1/trp1 | This study |
| FWY1556 | 1F, S2C | MAT A/α, Atg1(M_102_G)/Atg1(M_102_G), Ndt80::P_GAL1_-Ndt80:KanMX/Ndt80::P_GAL1_-Ndt80:KanMX, leu2::P_ACT1_-Gal4(848).ER:NatMX/leu2::P_ACT1_-Gal4(848).ER:NatMX, ∆rim4::Ura3/∆rim4::Ura3, his3::P_RIM4_-EGFP-Rim4(∆N424):His3/his3::P_RIM4_-EGFP-Rim4(∆N424):His3 | This study |
| FWY1967 | 1F | MAT A/α, Atg1(M_102_G)/Atg1(M_102_G), Ndt80::P_GAL1_-Ndt80:KanMX/Ndt80::P_GAL1_-Ndt80:KanMX, leu2::P_ACT1_-Gal4(848).ER:NatMX/leu2::P_ACT1_-Gal4(848).ER:NatMX, ∆rim4::Ura3/∆rim4::Ura3, his3::P_RIM4_-EGFP-Rim4(∆N424+KRm):His3/his3::P_RIM4_-EGFP-Rim4(∆N424+KRm):His3 | This study |
| FWY1563 | 1F | MAT A/α, Atg1(M_102_G)/Atg1(M_102_G), Ndt80::P_GAL1_-Ndt80:KanMX/Ndt80::P_GAL1_-Ndt80:KanMX, leu2::P_ACT1_-Gal4(848).ER:NatMX/leu2::P_ACT1_-Gal4(848).ER:NatMX, ∆rim4::Ura3/∆rim4::Ura3, his3::P_RIM4_-EGFP:His3/his3::P_RIM4_-EGFP:His3 | This study |
| FWY1471 | S1B, S2B, S2C, S6D, S6E | MAT A/α, Atg1(M_102_G)/Atg1(M_102_G), Ndt80::P_GAL1_-Ndt80:KanMX/Ndt80::P_GAL1_-Ndt80:KanMX, leu2::P_ACT1_-Gal4(848).ER:NatMX/leu2::P_ACT1_-Gal4(848).ER:NatMX, ∆rim4::Ura3/∆rim4::Ura3, his3::P_RIM4_-EGFP-Rim4:His3/his3::P_RIM4_-EGFP-Rim4:His3 | This study |
| FWY2312 | 1G | MAT A/α, Atg1(M_102_G)/Atg1(M_102_G), Ndt80::P_GAL1_-Ndt80:KanMX/Ndt80::P_GAL1_-Ndt80:KanMX, leu2::P_ACT1_-Gal4(848).ER:NatMX/leu2::P_ACT1_-Gal4(848).ER:NatMX, ∆rim4::Ura3/∆rim4::Ura3, his3::P_RIM4_-EGFP-Rim4(KRm):His3/his3, trp1::P_NUP49_-Nup49-mScarlet:Trp1/trp1 | This study |
| FWY2334 | 2B, 2C, 2D, 2E, 3J, 3K, 3L | MAT A/α, Atg1(M_102_G)/Atg1(M_102_G), Ndt80::P_GAL1_-Ndt80:KanMX/Ndt80::P_GAL1_-Ndt80:KanMX, leu2::P_ACT1_-Gal4(848).ER:NatMX/leu2::P_ACT1_-Gal4(848).ER:NatMX, ∆rim4::Ura3/∆rim4::Ura3, his3::P_RIM4_-EGFP-Rim4(FLm):His3/his3, trp1::P_NUP49_-Nup49-mScarlet:Trp1/trp1 | This study |
| FWY2316 | 2E | MAT A/α, Atg1(M_102_G)/Atg1(M_102_G), Ndt80::P_GAL1_-Ndt80:KanMX/Ndt80::P_GAL1_-Ndt80:KanMX, leu2::P_ACT1_-Gal4(848).ER:NatMX/leu2::P_ACT1_-Gal4(848).ER:NatMX, ∆rim4::Ura3/∆rim4::Ura3, his3::P_RIM4_-NES-EGFP-Rim4(FLm):His3/his3, trp1::P_NUP49_-Nup49-mScarlet:Trp1/trp1 | This study |
| FWY2043 | 2F | MAT A/α, Atg1(M_102_G)/Atg1(M_102_G), Ndt80::P_GAL1_-Ndt80:KanMX/Ndt80::P_GAL1_-Ndt80:KanMX, leu2::P_ACT1_-Gal4(848).ER:NatMX/leu2::P_ACT1_-Gal4(848).ER:NatMX, ∆rim4::Ura3/∆rim4::Ura3, his3::P_RIM4_-EGFP-Rim4(∆N424):His3/his3::P_RIM4_-mScarlet-Rim4:His3 | This study |
| FWY2117 | 2F | MAT A/α, Atg1(M_102_G)/Atg1(M_102_G), Ndt80::P_GAL1_-Ndt80:KanMX/Ndt80::P_GAL1_-Ndt80:KanMX, leu2::P_ACT1_-Gal4(848).ER:NatMX/leu2::P_ACT1_-Gal4(848).ER:NatMX, ∆rim4::Ura3/∆rim4::Ura3, his3::P_RIM4_-EGFP-Rim4(∆N424):His3/his3, trp1::P_PAB1_-mScarlet-Pab1:Trp1/trp1 | This study |
| FWY2143 | 2G, S2D | MAT A/α, Atg1(M_102_G)/Atg1(M_102_G), Ndt80::P_GAL1_-Ndt80:KanMX/Ndt80::P_GAL1_-Ndt80:KanMX, leu2::P_ACT1_-Gal4(848).ER:NatMX/leu2::P_ACT1_-Gal4(848).ER:NatMX, ∆rim4::Ura3/∆rim4::Ura3, his3::P_RIM4_-EGFP-Rim4:His3/his3 | This study |
| FWY2148 | 2G, S2D | MAT A/α, Atg1(M_102_G)/Atg1(M_102_G), Ndt80::P_GAL1_-Ndt80:KanMX/Ndt80::P_GAL1_-Ndt80:KanMX, leu2::P_ACT1_-Gal4(848).ER:NatMX/leu2::P_ACT1_-Gal4(848).ER:NatMX, ∆rim4::Ura3/∆rim4::Ura3, his3::P_RIM4_-EGFP-Rim4:His3/his3::P_RIM4_-EGFP-Rim4(∆N424):His3 | This study |
| FWY1557 | S2B | MAT A/α, Atg1(M_102_G)/Atg1(M_102_G), Ndt80::P_GAL1_-Ndt80:KanMX/Ndt80::P_GAL1_-Ndt80:KanMX, leu2::P_ACT1_-Gal4(848).ER:NatMX/leu2::P_ACT1_-Gal4(848).ER:NatMX, ∆rim4::Ura3/∆rim4::Ura3, his3::P_RIM4_-EGFP-Rim4(FLm):His3/his3::P_RIM4_-EGFP-Rim4(FLm):His3 | This study |
| FWY2061 | S2B | MAT A/α, Atg1(M_102_G)/Atg1(M_102_G), Ndt80::P_GAL1_-Ndt80:KanMX/Ndt80::P_GAL1_-Ndt80:KanMX, leu2::P_ACT1_-Gal4(848).ER:NatMX/leu2::P_ACT1_-Gal4(848).ER:NatMX, ∆rim4::Ura3/∆rim4::Ura3, his3::P_RIM4_-EGFP-Rim4(∆C289+FLm):His3/his3::P_RIM4_-EGFP-Rim4(∆C289+FLm):His3/ | This study |
| FWY2139 | S2C | MAT A/α, Atg1(M_102_G)/Atg1(M_102_G), Ndt80::P_GAL1_-Ndt80:KanMX/Ndt80::P_GAL1_-Ndt80:KanMX, leu2::P_ACT1_-Gal4(848).ER:NatMX/leu2::P_ACT1_-Gal4(848).ER:NatMX, ∆rim4::Ura3/∆rim4::Ura3, his3::P_RIM4_-NES-EGFP-Rim4(∆N424):His3/his3::P_RIM4_-NES-EGFP-Rim4(∆N424):His3 | This study |
| FWY1712 | 3A, S3A, S3B | MAT A/α, Atg1(M_102_G)/Atg1(M_102_G), Ndt80::P_GAL1_-Ndt80:KanMX/Ndt80::P_GAL1_-Ndt80:KanMX, leu2::P_ACT1_-Gal4(848).ER:NatMX/leu2::P_ACT1_-Gal4(848).ER:NatMX, ∆rim4::Ura3/∆rim4::Ura3, his3::P_RIM4_-EGFP-Rim4:His3/his3::P_CLB3_-CLB3-5’UTR-UbiY∆K-NanoLuciferase (CLB3 reporter):His3, trp1::P_CLB3_-CLB3-3×FLAG:Trp1/trp1 | This study |
| FWY2042 | 3B | MAT A/α, Atg1(M_102_G)/Atg1(M_102_G), Ndt80::P_GAL1_-Ndt80:KanMX/Ndt80::P_GAL1_-Ndt80:KanMX, ura3::P_TDH3_-Gal4(848).ER:Ura3/ura3::P_TDH3_-Gal4(848).ER:Ura3, Rim4-3×V5:His3/Rim4-3×V5:His3, Pab1-3×FLAG:NatMX/Pab1-3×FLAG:NatMX, | This study |
| FWY1184 | 3F, 3G, 3H, 4G, S5B, S5C, S5D | MAT A/α, Atg1(M_102_G)/Atg1(M_102_G), Ndt80::P_GAL1_-Ndt80:KanMX/Ndt80::P_GAL1_-Ndt80:KanMX, ura3::P_TDH3_-Gal4(848).ER:Ura3/ura3::P_TDH3_-Gal4(848).ER:Ura3, Rim4::P_RIM4_-EGFP-Rim4:Rim4/Rim4, his3::P_ATG8_-mScarlet-Atg8:His3/his3::P_ATG8_-mScarlet-Atg8:His3 | This study |
| FWY2398 | S3C, S3D | MAT A/α, Atg1(M_102_G)/Atg1(M_102_G), Ndt80::P_GAL1_-Ndt80:KanMX/Ndt80::P_GAL1_-Ndt80:KanMX, ura3::P_TDH3_-Gal4(848).ER:Ura3/ura3::P_TDH3_-Gal4(848).ER:Ura3, leu2::P_PAB1_-EGFP-Pab1:Leu2/leu2::P_PAB1_-EGFP-Pab1:Leu2, his3::P_ATG8_-mScarlet-Atg8:His3/his3 | This study |
| FWY1598 | 4A, 4B, S4C | MAT A/α, Atg1(M_102_G)/Atg1(M_102_G), Ndt80::P_GAL1_-Ndt80:KanMX/Ndt80::P_GAL1_-Ndt80:KanMX, ura3::P_TDH3_-Gal4(848).ER:Ura3/ura3, leu2::P_ACT1_-Gal4(848).ER:NatMX/leu2, ∆rim4::Ura3/∆rim4::mCherry-GFP, his3::P_RIM4_-EGFP-Rim4:His3/his3 | This study |
| FWY1824 | 4C, 4D, S4C | MAT A/α, Atg1(M_102_G)/Atg1(M_102_G), Ndt80::P_GAL1_-Ndt80:KanMX/Ndt80::P_GAL1_-Ndt80:KanMX, ura3::P_TDH3_-Gal4(848).ER:Ura3/ura3, leu2::P_ACT1_-Gal4(848).ER:NatMX/leu2, ∆rim4::Ura3/∆rim4::mCherry-GFP, his3::P_RIM4_-EGFP-Rim4:His3/his3, ∆atg11::Leu2/∆atg11::Leu2 | This study |
| FWY1904 | S4A, S4B | MAT A/α, Atg1(M_102_G)/Atg1(M_102_G), Ndt80::P_GAL1_-Ndt80:KanMX/Ndt80::P_GAL1_-Ndt80:KanMX, ura3::P_TDH3_-Gal4(848).ER:Ura3/ura3, leu2::P_ACT1_-Gal4(848).ER:NatMX/leu2, ∆rim4::Ura3/∆rim4::Ape1-GFP, his3::P_RIM4_-EGFP-Rim4:His3/his3 | This study |
| FWY1819 | 4E, 4F, 4G | MAT A/α, Atg1(M_102_G)/Atg1(M_102_G), Ndt80::P_GAL1_-Ndt80:KanMX/Ndt80::P_GAL1_-Ndt80:KanMX, leu2::P_ACT1_-Gal4(848).ER:NatMX/leu2::P_ACT1_-Gal4(848).ER:NatMX, ∆rim4::Ura3/∆rim4::Ura3, ∆atg11::Leu2/∆atg11::Leu2, his3::P_RIM4_-EGFP-Rim4:His3/his3::P_ATG8_-mScarlet-Atg8:His3 | This study |
| FWY643 | 5B, 5D, 5E | MAT A/α, Atg1(M_102_G)/Atg1(M_102_G), Ndt80::P_GAL1_-Ndt80:Trp1/Ndt80::P_GAL1_-Ndt80:Trp1, ura3::P_TDH3_-GAL4(848).ER:Ura3/ura3::P_TDH3_-GAL4(848).ER:Ura3 | (Feng et al., 2022) |
| FWY2407 | 5C | MAT A/α, FLAG-Atg1(M_102_G)/FLAG-Atg1(M_102_G), Ndt80::P_GAL1_-Ndt80:Trp1/Ndt80::P_GAL1_-Ndt80:Trp1, Ura3::P_TDH3_-GAL4(848).ER:Ura3/ura3::P_TDH3_-GAL4(848).ER:Ura3 | (Feng et al., 2022) |
| FWY1621 | S3A, 6A-6H, S6B, S6C, S6F | MAT A/α, Atg1(M_102_G)/Atg1(M_102_G), leu2::P_ACT1_-Zif268-ER-VP14(ZEV)+P_ZEV_-TEV:Leu2/leu2::P_ACT1_-Zif268-ER-VP14(ZEV)+P_ZEV_-TEV:Leu2, N-Deg-Rim4-3×V5/N-Deg-Rim4, his3::P_CLB3_-CLB3-5’UTR-UbiY∆K-NanoLuciferase:His3/his3 | This study |
| FWY516 | S6A, S6B | MAT A/α, Atg1(M_102_G)/Atg1(M_102_G), Rim4-3×V5:His3/Rim4-3×V5:His3 | This study |
| FWY473 | 6A, S6A, S6B | MAT A/α, Atg1(M_102_G)/Atg1(M_102_G), leu2::P_ACT1_-Zif268-ER-VP14(ZEV)+P_ZEV_-TEV:Leu2/leu2::P_ACT1_-Zif268-ER-VP14(ZEV)+P_ZEV_-TEV:Leu2, N-Deg-Rim4-3×V5/N-Deg-Rim4-3×V5 | This study |
