## Supplementary material for "Autophagy inspects Rim4-mRNA interaction to safeguard programmed meiotic translation": Table S2

**Table S2**. Plasmids used in this study

| Plasmid Name | Description | Source |
| --- | --- | --- |
| pFW1 | pRS304>P_TDH3_-Gal4(848).ER:Ura3 | (Wang et al., 2020) |
| pFW127 | pAGL>P_ACT1_-Gal4(848).ER:NatMX | This study |
| pFW18 | pKL-URA>Ura3 | A gift from Denic Lab |
| pFW52 | pKT0128>yeGFP:His3 | A gift from Denic Lab |
| pFW72 | pET29b>Rim4-6×His | This study |
| pFW95 | pET29b>Rim4-EGFP-6×His | This study |
| pFW353 | pET29b>Rim4(FLm)-6×His | This study |
| pFW81 | pET29b>Rim4(ΔC289)-6×His | This study |
| pFW394 | pET28a>6×His-3×FLAG-Cdc14 | This study |
| pFW103 | pET29b>Pab1-6×His | This study |
| pFW115 | pET29b>P_T7_-CLB3-5’UTR-Clb3-CLB3-3’UTR | This study |
| pFW43 | pNH605>P_ACT1_-Zif268-ER-VP16+P_ZEV_-TEV | This study |
| pFW46 | pNH605> P_ACT1_-Zif268-ER-VP16+P_ZEV_-N-Deg-Rim4 |  |
| pFW338 | pRS304>P_NUP49_-Nup49-mScarlet:Trp1 | This study |
| pFW337 | pRS304>P_PAB1_-mScarlet-Pab1:Trp1 | This study |
| pFW289 | pRS303>P_RIM4_- mScarlet-Rim4:Trp1 | This study |
| pFW281 | pRS303>P_ATG8_-mScarlet-Atg8:His3 | This study |
| pFW208 | pRS303>P_RIM4_-EGFP-Rim4:His3 | This study |
| pFW250 | pRS303>P_RIM4_-EGFP-Rim4(FLm):His3 | This study |
| pFW297 | pRS303> P_RIM4_-EGFP-Rim4(KRm):His3 | This study |
| pFW290 | pRS303>P_RIM4_-EGFP-Rim4(∆C289):His3 | This study |
| pFW249 | pRS303>P_RIM4_-EGFP-Rim4(∆N424):His3 | This study |
| pFW313 | pRS303>P_RIM4_-EGFP-Rim4(∆N424+KRm):His3 | This study |
| pFW256 | pRS303>P_RIM4_-EGFP:His3 | This study |
| pFW347 | pRS303>P_RIM4_-NES-EGFP-Rim4(FLm):His3 | This study |
| pFW348 | pRS303>P_RIM4_-NES-EGFP-Rim4(∆N424):His3 | This study |
| pFW324 | pRS303>P_RIM4_-EGFP-Rim4(∆C289+FLm):His3 | This study |
| pFW225 | pRS303>P_CLB3_-CLB3-5’UTR-UbiY∆K-NanoLuciferase: His3 | This study |
| pFW342 | pRS304>P_CLB3_-CLB3-5’UTR-CLB3-3×FLAG:Trp1 | This study |
| pFW398 | pRS305>P_PAB1_-EGFP-Pab1:Leu2 | This study |
