## Supplementary material for "Autophagy inspects Rim4-mRNA interaction to safeguard programmed meiotic translation": Table S3

**Table S3**. Oligos used in this study

| Oligo Number | Oligo Name | Sequence (5’ -> 3’) |
| --- | --- | --- |
| qPCR primers | | |
| oFW1392 | ACT1-F | AGAGTTGCCCCAGAAGAACA |
| oFW1393 | ACT1-R | GGCTTGGATGGAAACGTAGA |
| oFW1496 | CLB3-F | tggtctccgctcaacctagt |
| oFW1497 | CLB3-R | cggaagaatttgttcttgtg |
| oFW1454 | Luc-F | GGTGTCCGTAACTCCGATCC |
| oFW1455 | Luc-R | TCGATCTGGCCCATTTGGTC |
| oFW929 | RIM4-F | atcacgcaaaactttaac |
| oFW287 | RIM4-R | ctaccacttccacatccact |
| oFW1458 | SPS1-F | GGTCAAGAACCCCTCCATCG |
| oFW1459 | SPS1-R | TCGGAATGCTCCAGGTTGAC |
| oFW1460 | YSP2-F | CACTTCTGGTCCAGCCTCTG |
| oFW1461 | YSP2-R | TGAAAGGGGCGTAGTCGATG |
| oFW1462 | YFL012W-F | AGGACCATTGCGTCTTCCTC |
| oFW1463 | YFL012W-R | TCGTTTCATCCGTTGTTGCG |
| oFW1464 | SPO20-F | caaggggtggaaagacagcg |
| oFW1465 | SPO20-R | TTCAAGTCACCACTCACCGC |
| oFW1456 | AMA1-F | GCACAGCAAGTCTGTGGTATT |
| oFW1457 | AMA1-R | CACAGCCGCTTTATGTGGAA |
| oFW1466 | YDR042C-F | ctgaccagaagtggaattcg |
| oFW1467 | YDR042C-R | ctattttttagtccatcggc |
| oFW1468 | CTS2-F | GGTAGGACACTCAGCTCAGC |
| oFW1469 | CTS2-R | gatgtccgccattgttaaag |
| oFW1470 | SPR6-F | CTTGTCAGGACCCGTGACTC |
| oFW1471 | SPR6-R | TGCTGACATGCACACCTCTC |
| oFW1472 | YGL230C-F | TCGCTTGCTCTTTTATTTGTGCT |
| oFW1473 | YGL230C-R | TCGCATATGGAACGCATTGC |
| oFW1474 | SGA1-F | AGAACGGTCGTCTCCCTTTG |
| oFW1475 | SGA1-R | TATGTGTCGGCAATCATGGG |
| oFW1476 | CUZ1-F | TGCTAAAGGTGACCCCAAGA |
| oFW1477 | CUZ1-R | CCTACCCACAGGCCACATTT |
| oFW1478 | SPR2-F | ACAGAAGAAAGTCGGGGCAG |
| oFW1479 | SPR2-R | TGTTGCAGTACTTGTCGGCA |
| oFW1480 | SSP2-F | ACGGCAGGTTCTCAGAATCC |
| oFW1481 | SSP2-R | AACCTCCGCGTACTTGAGAC |
| oFW1482 | SPS4-F | CAGCTCAGGGATACGACAGC |
| oFW1483 | SPS4-R | AGGCTTGCCCTTTGTGTCTA |
| oFW472 | ATG1-F | CAACGAGATTGTTTGGTG |
| oFW473 | ATG1-R | TGCTGTAGGTGATCTATTC |
| oFW478 | ATG8-F | GTCTGAATATCCATTTGAAA |
| oFW479 | ATG8-R | CAGGTATCCTATTCTTGAA |
| oFW1484 | CDC5-F | TACACCATATGCGGAACACC |
| oFW1485 | CDC5-R | TGCTTGGAAGGGTGGCTTAC |
| oFW1486 | CLB4-F | AAATAGGTTGGCCAGGACCC |
| oFW1487 | CLB4-R | CTCAGAAAATACGCGCCAGC |
| oFW1488 | SPO12-F | TTGCATCACCGACTGACAGG |
| oFW1489 | SPO12-R | cttcatcgatttctacatcc |
| oFW1490 | PGK1-F | GGTCCACCAGGTGTTTTCGA |
| oFW1491 | PGK1-R | acaccgtacttcttagcgac |
| oFW1492 | HXK1-F | ACCCAGGTTTCAAGGAAGCC |
| oFW1493 | HXK1-R | TGCACCTGAACCATCCTCAG |
| oFW1436 | POR1-F | gccggtttgcttgtcattca |
| oFW1437 | POR1-R | ctaccccagctgcctttgat |
| oFW1439 | RPL13A-F | TAGAGCTACCAGAGCCGCTA |
| oFW1440 | RPL13A-R | TAGCTCTTTCAGCCAAGGTG |
| oFW1442 | RPS1B-F | gcttccgatgctttgaaggg |
| oFW1443 | RPS1B-R | ccgtggaagttggtcaacaag |
| EMSA probe | | |
| oFW1405 | CLB3 5UTR | Ttttagctgcttgcttagaagaaaagaaaagaaaat  TgcagagaaaaagctatttattgtttctgtccttcgC  tttaaaacatagagatattttgtcctttttatcattgc  ttattataa |
